# Aging disrupts transcriptional programs for memory updating in the dorsal hippocampus and reveals a required role for *Tent5a* in reconsolidation

**DOI:** 10.64898/2026.09.08.750076

**Authors:** Chad A. Brunswick, Annie G. Defina, Trinity A. Wood, Aswathy Sebastian, Alexandria R. McKenna, Shoko Murakami, Pu-Hsun Chiu, Shannon M. Jordan, Gretchen C. Pifer, Istvan Albert, Janine L. Kwapis

## Abstract

Existing memories can be updated with new information through a process known as reconsolidation, though the molecular mechanisms underlying this process are poorly understood. Additionally, memory updating is impaired with aging, but little is known about how this impairment occurs. Here, we used a hippocampus-dependent memory updating task alongside transcriptomics to identify which genes are upregulated in the dorsal hippocampus specifically during a memory update in both young adult mice, which show successful memory updating, and in old mice, which do not. In young mice, we observed that distinct transcriptional programs were activated by reconsolidation-dependent memory updating and memory retrieval without new information. In old mice, similar transcriptional programs were engaged by both memory updating and memory retrieval. From our sequencing results, we identified *Tent5a* as a novel regulator of reconsolidation-mediated memory updating, and we demonstrate that hippocampal expression of *Tent5a* is necessary for this process. Together, these results expand our understanding of both the underlying transcriptional mechanisms of memory updating and how these mechanisms go awry in the aged brain.

**Significance Statement:** Many studies have examined the transcriptional mechanisms that contribute to memory formation, typically relying on highly controlled exposures to isolated stimuli. However, memories are not formed in isolation, and the brain constantly integrates new experiences into existing memories and information stores to drive optimal behavior. This process of memory updating is preferentially impaired with aging yet has received relatively little attention in the scientific literature. Here we compared gene expression during memory updating in the dorsal hippocampus of young adult mice (which successfully update memories) and old mice (which do not). These results expand our understanding of how transcriptional mechanisms change with age and how these changes contribute to age-related deficits in understudied aspects of cognition, like memory updating.

## Introduction

Memories can be updated to reflect new or changed information (see Brunswick et al., 2025; Wahlheim and Zacks, 2025) through a process known as reconsolidation. During reconsolidation, memories are transiently destabilized by protein degradation before being subsequently restabilized through protein synthesis (see Tronson and Taylor, 2007; Bellfy and Kwapis, 2020; Jardine et al., 2022). As a result of this destabilization, memories undergoing reconsolidation are susceptible to the effects of amnestic agents (e.g., the protein synthesis inhibitor anisomycin (Nader et al., 2000)) and also capable of incorporating new information (Monfils et al., 2009; Gräff et al., 2014; Jarome et al., 2015; Kwapis et al., 2017, 2019a; Ferrara et al., 2019), i.e., updating. However, the mechanisms underlying this process are poorly understood. Both initial memory consolidation (Flexner et al., 1963; Davis and Squire, 1984) and subsequent reconsolidation (Nader et al., 2000; Sangha et al., 2003; Duvarci et al., 2008) are dependent on gene expression and protein synthesis. A myriad of studies have explored the transcriptional mechanisms necessary for memory consolidation (e.g., Jones et al., 2001; Stork et al., 2001; Mei et al., 2005; Kwapis et al., 2018; Bellfy et al., 2023), but far fewer have examined which genes are required for memory reconsolidation. Observations that reconsolidation disruption can be reversed following either the passage of time (Lattal and Abel, 2004) or a reminder of the initial training (Trent et al., 2015) suggest that reconsolidation is not a reiteration of initial memory consolidation but rather a fundamentally distinct process that is capable of updating established memories with new information.

Recent reports indicate that memory updating may be more susceptible to the effects of aging than initial memory formation (Kwapis et al., 2019a; Amelchenko et al., 2023; Mau et al., 2023; Jardine et al., 2025). This further suggests that mechanistic differences may exist between consolidation and reconsolidation and also highlights the importance of understanding these mechanisms to potentially improve cognition in old age. Nonetheless, the transcriptional mechanisms of reconsolidation-dependent memory updating remain largely unknown (see Tronson and Taylor, 2007; Bellfy and Kwapis, 2020), and no investigation to date has examined how these mechanisms change as a result of aging. To address this, we used a memory updating task to investigate hippocampal gene expression following reconsolidation-dependent memory updating in young adult and old mice.

## Materials and Methods

### Subjects

All mice used for these experiments were C57BL/6J. Young adult mice were between 3 and 4 months of age at the start of experiments and were acquired from the Jackson Laboratory (Bar Harbor, ME) while old mice were between 18 and 20 months of age at the start of experiments and were acquired from the NIA Aged Rodent Colony (maintained by Charles River, Raleigh, NC). Mice were housed in groups of four in a temperature- and humidity-controlled environment with *ad libitum* access to food and water and a 12-hour light/dark cycle. All experiments were performed during the light phase of the cycle, as this is when we find C57BL/6J mice learn spatial memory tasks best (Bellfy et al., 2023). Mice were randomly assigned to groups and appropriately counterbalanced. All experiments were approved by the Institutional Animal Care and Use Committee at the Pennsylvania State University.

### Objects in Updated Locations (OUL)

OUL was performed as previously described (Kwapis et al., 2019a; Wright et al., 2020). In brief, mice were handled for 4 consecutive days in the behavior room (2 min each day) and then habituated to the polypropylene arenas (23.0 cm x 30.0 cm x 23.0 cm) for 5 min each day for 6 consecutive days. Following habituation, mice underwent 3 consecutive days of 10-min training in which identical objects (glass 200mL tall-form beakers filled with hydraulic cement) were presented in locations A1 and A2 of the arena. This 3-day training protocol allows old mice to successfully learn the training prior to the update (Kwapis et al., 2019a) and was used for both age groups to maintain consistency. Twenty-four hours following the last training, mice underwent the 5-min update session with objects presented in locations A1 and A3. Finally, mice were given a 5-min retention test the next day in which objects were presented in all previously seen locations (A1, A2, and A3) and in the novel location A4. The orientations of these locations were counterbalanced as previously described (Wright et al., 2020).

Movement data were calculated using Ethovision (Noldus, Leesburg, VA) to verify that mice habituated to the OUL context by exhibiting a decrease in distance moved prior to beginning the training sessions. Object investigation was determined by manually scoring behavioral videos on a frame-by-frame basis by a blinded investigator using the GUI of DeepEthoGram (Bohnslav et al., 2021). Investigation was defined as any frame in which the mouse had all four paws on the ground with its nose pointed directly at the object and within 1 cm of the object and was not otherwise biting, climbing, digging, or rearing on the object (Vogel-Ciernia and Wood, 2014). Discrimination indices (DIs) were calculated to quantify memory using the general formula DI = (t_n_ – t_f_)/(t_n_ + t_f_) × 100% where t_n_ is the time spent investigating an object in a novel location and t_f_ is the time spent investigating an object in a familiar location. For the OUL test session, three separate DIs were calculated per animal, each comparing the novel location A4 against a different familiar location (A1, A2, or A3). Preference for A4 over A1 or A2 was interpreted as memory for the initial training, while preference for A4 over A3 was interpreted as memory for the update. DIs were additionally calculated during each training session, to verify animals exhibited no innate preference for a particular side of the arena, and during the update session, to confirm animals successfully learned the initial training. To minimize the likelihood of misinterpreting animal behavior in our task, we maintain strict exclusion criteria. We exclude from behavioral analyses any animal that fails to explore all four objects during the test session, investigates for less than three seconds total during the test session, exhibits a DI greater than 20 or less than −20 on all three training days, or exhibits a DI less than −20 during the update session, though no animals met these exclusion criteria in this study.

### RNA Sequencing

Animals were euthanized via cervical dislocation 60 min after behavior and decapitated with surgical scissors. Brains were removed from the skull and flash-frozen in 2-methylbutane (Fisher Scientific, Waltham, MA). Brains were stored at −80 °C before being sectioned with a Leica CM1950 Cryostat (Leica Biosystems, Wetzlar, Germany). Circular punches 1 mm in diameter and 500 μm thick were collected from the dorsal hippocampus and stored at −80 °C prior to RNA extraction via RNeasy Mini Kits (Qiagen, Germantown, MD). Total RNA quality and concentration was assessed using an RNA Screen Tape on a TapeStation 4150 (Agilent, Santa Clara, CA). A uniquely dual indexed cDNA library was prepared by the Penn State Genomics Core using the Illumina Stranded mRNA library prep kit (Illumina). Library quality and size was assessed with the DNA 5000 Screen Tape on the TapeStation 4150, and concentration was determined via qPCR using the KAPA Library Quantification Kit for Illumina platforms (Kapa Biosystems, Wilmington, MA). A 10 nM equimolar pool of all libraries was made, and the balance was confirmed by 150 x 150 paired-end sequencing on the Illumina MiSeq Nano, followed by sequencing on the NextSeq 2000 using a P3 kit (Illumina, San Diego, CA) with single-read 100 nt sequencing. Single-end 100 bp RNA-seq reads were generated from young and old mouse samples across multiple conditions. Read quality was assessed using FastQC. Reads were aligned to the mouse reference genome mm10 using HISAT2 with default parameters. Gene expression was quantified using Salmon directly from FASTQ files, and transcript-level estimates were summarized to the gene level. Differential expression analysis was performed in R using the DESeq2 package, while accounting for unwanted variation with RUVSeq (Risso et al., 2014). Genes showing no evidence of differential expression across any condition in a first-pass DESeq2 analysis without unwanted variation correction were used as empirical negative controls to estimate unwanted variation factors using the RUVg method. The estimated unwanted variation factors were included in the DESeq2 design formula to identify significantly differentially expressed genes.

### siRNA Administration

For the knockdown experiment, Accell SMARTpool small interfering RNA (siRNAs; Dharmacon, Lafayette, CO) were diluted in ddH_2_O to a final total concentration of 10μM and administered bilaterally to the DH via stereotaxic surgery as previously described (López et al., 2016; Kwapis et al., 2018). Briefly, mice were anesthetized with isoflurane in oxygen (induced at 3%, maintained at 1-2%) and placed in the stereotaxic head frame, and then a craniotomy was performed with a micromotor drill. Injection needles were lowered at a rate of 0.2 mm/15 sec to the final coordinates (from Bregma) of AP: −2.00 mm; ML: ±1.50 mm; and DV: −1.50 mm. After a 2-min pause, 1 μL of siRNA was infused at a rate of 15 μL/hr. The injectors were left in place for 5 min after the completion of the infusion, after which they were raised by 0.1 mm and allowed to rest for an additional 2 min. Finally, injectors were withdrawn from the injection site at a rate of 0.1 mm/15 sec. Animals were administered 5 mg/kg subcutaneous meloxicam as an analgesic, given 0.5 mL sterile saline for hydration, and allowed to recover in a clean, heated cage. Mice were given a full day of recovery, and behavior resumed 48 hrs after surgery to ensure maximal knockdown.

### Experimental Design and Statistical Analyses

Data are represented as Mean ± SEM in all figures. Sample sizes were chosen on the basis of previous studies using similar molecular and behavioral assays (e.g., Kwapis et al., 2018, 2019a; Brunswick et al., 2023), but no statistical method was used to predetermine sample size. All analyses were conducted in GraphPad Prism 10 (GraphPad Software, San Diego, CA). Data normality was verified with the Shapiro-Wilk test and variances between groups were checked with an F test prior to subsequent analysis. Data were analyzed using one-sample t-tests (**Fig. 1B, 5B, 5D, 5E, 5F**), unpaired Student’s t-tests (**Fig. 5B, 5C, 5D, 5E, 5F, 5G**), two-way ANOVAs (**Fig. 1B, 1C**), two-way repeated-measures ANOVAs (**Fig. S3**), three-way repeated-measures ANOVAs (**Fig. S1**), mixed effect analyses (**Fig. S1**), or basic linear regression (**Fig. 2C, 2D**). For all statistical tests, α was set to 0.05 and all comparisons were two-tailed. Any animal that exhibited a test-session DI more than 2 standard deviations away from their group mean was excluded from all analyses (4 mice across all experiments).

**Figure 1.**
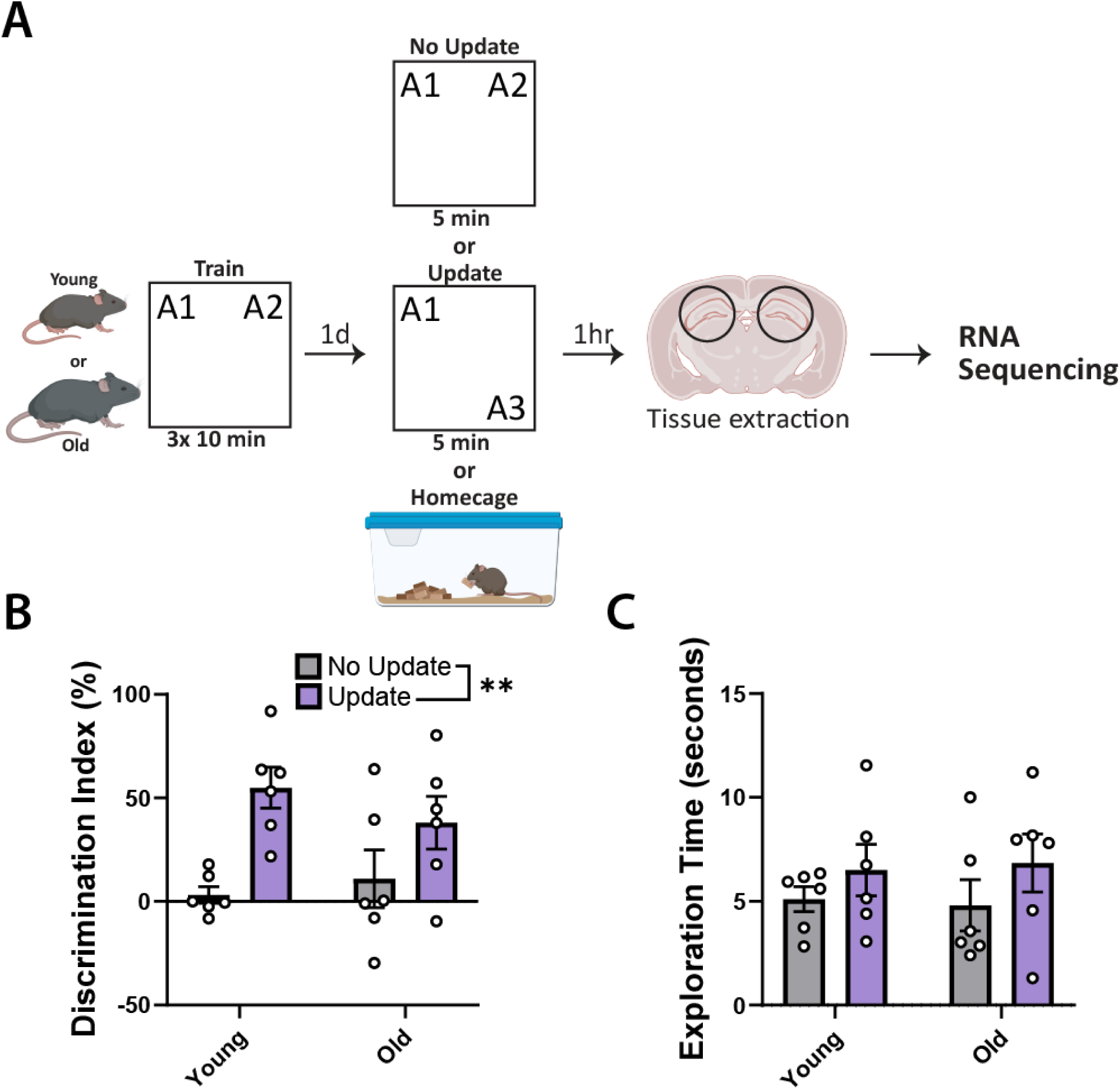
Experimental design and behavioral data for RNA sequencing experiment. **A.** Experimental schematic. Young and old mice were given three consecutive training days to form an initial memory for positions A1 and A2. Twenty-four hours later, mice were either given a repeat of the training experience (No Update), placed back into the behavioral arena but with an object present in the update position A3 (Update) or left undisturbed in their homecage. An hour after this experience, mice were sacrificed and hippocampal punches were taken for RNA-sequencing. **B.** Discrimination indices during the update day. Mice in the U group demonstrated intact memory for the initial training (one-sample t-tests against 0: young NU: t_5_ = 0.79, p = 0.46; young U: t_5_ = 5.57, p = 0.0026; old NU: t_5_ = 0.78, p = 0.47; old U: t_5_ = 2.99, p = 0.031) and also exhibited significantly higher discrimination indices than mice in the NU group with no differences between age groups (two-way ANOVA, effect of age: F_(1,20)_ = 0.18, p = 0.68; effect of behavior: F_(1, 20)_ = 13.20, p = 0.0017; interaction: F_(1,20)_ = 1.28, p = 0.27). **C.** Total exploration time during the update day. No significant differences in exploration were observed between age or behavioral groups (two-way ANOVA, effect of age: F_(1,20)_ = 0.00037, p = 0.98; effect of behavior: F_(1, 20)_ = 2.18, p = 0.16; interaction: F_(1,20)_ = 0.075, p = 0.79). All data are presented as mean ± SEM. n = 6, 6, 6, 6; all male.

**Figure 2.**
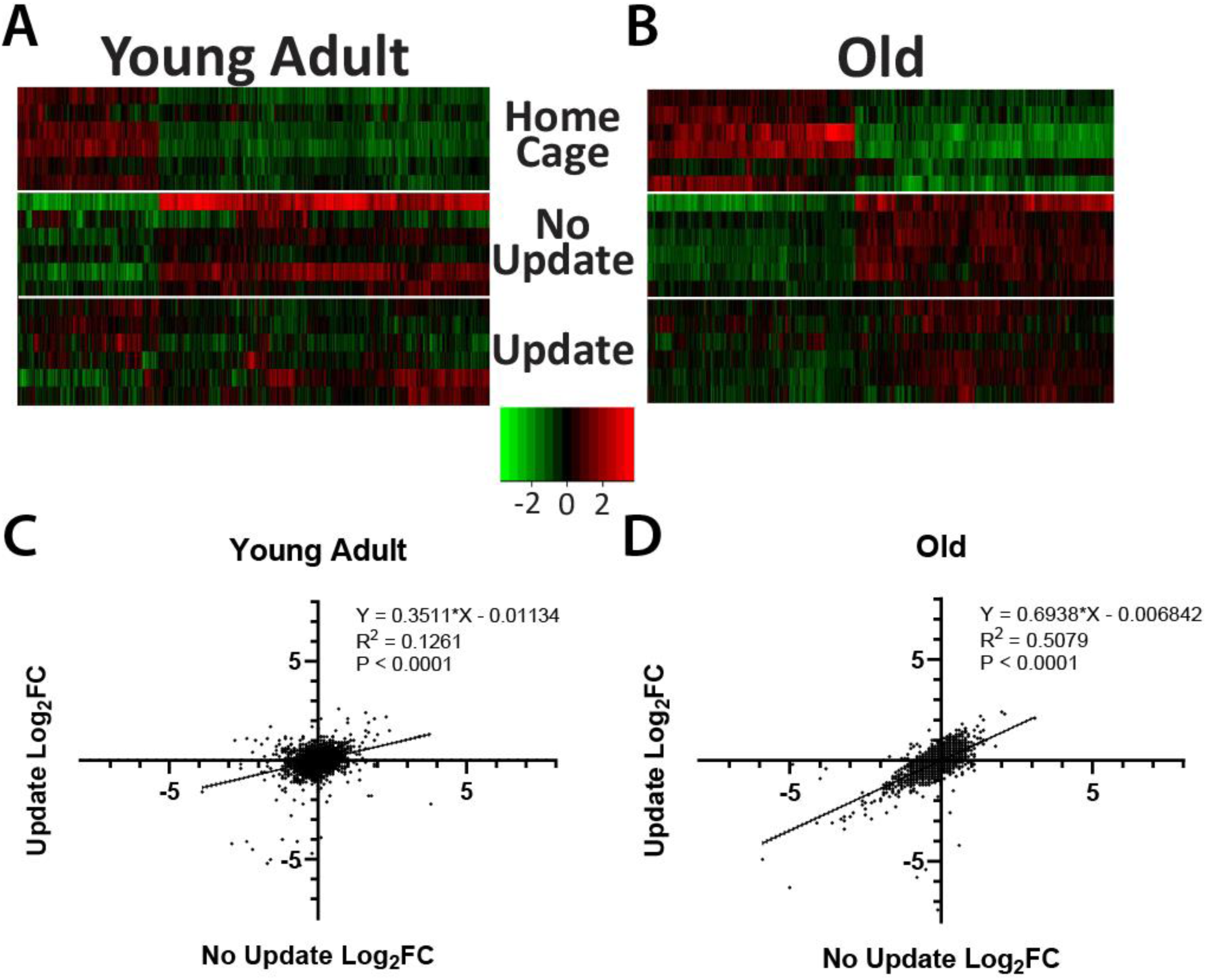
Both memory updating and memory retrieval drive large changes in hippocampal gene expression in young and old mice. A &. **B.** Heatmaps comparing hippocampal gene expression following the no update or update experience in young (**A**) and old mice (**B**). Each column represents an individual gene with red denoting upregulation and green denoting downregulation. **C.** Line plot comparing the Log_2_FC in the young update and no update groups for all detected genes (simple linear regression, R^2^ = 0.13, F_(1, 15398)_ = 2221, p < 0.0001, Y = 0.35X – 0.011). **D.** Same as **C** but comparing Log_2_FC in the old animals (simple linear regression, R^2^ = 0.51, F_(1, 15434)_ = 15930, p < 0.0001, Y = 0.69X - 0.0068).

### Data Availability

The data supporting the findings of this study are uploaded to the Inter-university Consortium for Political and Social Research (ICPSR). Additional data files and materials are available from the corresponding author upon reasonable request. RNA sequencing data have been deposited in the NCBI’s Gene Expression Omnibus and are accessible through GEO accession number GSE342530.

## Results

### Hippocampal gene expression during memory updating is disrupted by aging

In order to explore gene expression during reconsolidation-mediated updating, we used the Objects in Updated Locations (OUL) memory updating task (Wright et al., 2020). OUL is a hippocampus-dependent incidental learning task in which mice initially learn the positions of two identical objects (deemed A1 and A2). Following this training, mice undergo an update session in which they are presented with one object in a previously seen position (A1) and one in a novel update position A3. Finally, mice are tested on both their memory for the initial training and the update session. We have demonstrated that this update session is a memory update rather than a separate memory, as it relies on largely the same hippocampal neurons as the initial training (Kwapis et al., 2019a), at least in young mice (Brunswick et al., 2026). Further, we have shown that administration of the protein synthesis inhibitor anisomycin immediately following the update session interrupts memory for both the update and the initial training in young mice (Kwapis et al., 2019a), consistent with the expected effects of interrupting reconsolidation. Finally, we have also demonstrated that old mice exhibit updating impairments in OUL even when the initial training is successfully learned (Kwapis et al., 2019a). Thus, this task is ideal for examining the transcriptional mechanisms underlying reconsolidation-mediated memory updating and investigating how these mechanisms change in old age, when memory updating is impaired.

To assess hippocampal gene expression during memory updating, young adult and old mice were split into 3 groups: homecage (HC), no-update (NU), and update (U). All 3 groups underwent the handling, habituation, and training days of OUL. On the update day, the HC group was left undisturbed in their homecages in the colony room, while the NU group received a no-update session consisting of a 5 min repeat of the training positions (A1 and A2) without exposure to the updated position A3, and the U group underwent the typical 5 min OUL update, with one object in a familiar position (A1) and another object in the updated position A3. One hour after this experience, these animals were sacrificed (counterbalanced with HC mice), and RNA from hippocampal punches was extracted for sequencing (**Fig. 1A**).

During habituation, mice acclimated to the arena as expected (**Fig. S1A**), with old animals moving significantly less, consistent with age-related decreases in locomotion. During the training sessions, mice behaved as expected, demonstrating no preference for either side of the arena (**Fig. S1B**) and exhibiting decreasing exploration time across sessions (**Fig. S1C**).

On the update day, mice in the U groups demonstrated preference for position A3 over A1, indicating intact memory for the initial training (**Fig. 1B**; one-sample t-tests of update session DIs against 0: young NU: t_5_ = 0.79, p = 0.46; young U: t_5_ = 5.57, p = 0.0026; old NU: t_5_ = 0.78, p = 0.47; Old U: t_5_ = 2.99, p = 0.031), and these groups demonstrated significantly higher DIs than mice in the NU groups (two-way ANOVA, effect of age: F_(1,20)_ = 0.18, p = 0.68; effect of behavior: F_(1, 20)_ = 13.20, p = 0.0017; interaction: F_(1,20)_ = 1.28, p = 0.27). Additionally, we observed no differences in total investigation time across our groups (**Fig. 1C**; two-way ANOVA, effect of age: F_(1,20)_ = 0.00037, p = 0.98; effect of behavior: F_(1, 20)_ = 2.18, p = 0.16; interaction: F_(1,20)_ = 0.075, p = 0.79). Although gene expression is required for memory consolidation and reconsolidation, many other forms of stimulation, including memory retrieval (Peixoto et al., 2015), can drive DH gene expression. Thus, we can identify updating-specific genes by comparing the differentially expressed genes (DEGs) in the young U group with those identified in the young NU group.

Initially, we compared broad patterns of gene expression across each of our three behavioral groups (HC, NU, and U). In young mice, we noted largely distinct gene expression profiles for each group (**Fig. 2A**), suggesting that the young DH might initiate distinct transcriptional programs based on whether new information is present or absent during memory retrieval. In old mice, however, we observed more overall similarities between the gene expression profile of the NU group and the U group (**Fig. 2B**), indicating the old DH exhibits less transcriptional differentiation between memory retrieval and memory updating. From here, we sought to identify genes altered in response to either the NU or U condition in each age group using 4 total comparisons: young NU vs young HC, young U vs young HC, old NU vs old HC, and old U vs old HC. Genes that exhibited significant differences (FDR < 0.05) in any of these comparisons were identified to produce 4 unique list of DEGs: young NU DEGs (891 genes; **Table S1**), young U DEGs (433 genes; **Table S1**), old NU DEGs (1572 genes; **Table S2**), and old U DEGs (1226 genes; **Table S2**). Next, we compared, for all detected genes, the change in expression observed in the NU vs HC comparison with that observed in the U vs HC comparison separately for young adult and old mice. In the young adult DH, we observed a significant correlation between the NU and U log_2_ fold changes with a slight slope (**Fig. 2C**; simple linear regression, R^2^ = 0.13, F_(1, 15398)_ = 2221, p < 0.0001, Y = 0.35X – 0.011), while in the old DH the NU and U log_2_ fold changes showed a much greater degree of similarity, with both a stronger correlation and a slope closer to 1.0 (**Fig. 2D**; simple linear regression, R^2^ = 0.51, F_(1, 15434)_ = 15930, p < 0.0001, Y = 0.69X - 0.0068). Together, these data indicate that the young DH engages behavior-specific patterns of transcription while the old DH relies more on a shared transcriptional program across these different behaviors.

### Identification of update-specific genes in the young DH

As the molecular mechanisms underlying memory updating remain poorly understood, we were interested in identifying a putative transcriptional program for reconsolidation-mediated memory updating in our young animals. To identify genes involved in successful updating, we cross-referenced the upregulated DEGs identified in the young NU group (**Fig. 3A**) with those from the young U group (**Fig. 3B**). We reasoned that reconsolidation-specific genes would be unique to the U group, as the NU group receives no new information during the final session and we have shown that this experience does not initiate reconsolidation (Kwapis et al., 2019a). We identified 625 upregulated DEGs in the young NU group and 284 DEGs in the young U group, with 86 of these DEGs common to both groups (**Fig. 3C**). The 198 genes that were upregulated in young mice uniquely in the U group therefore represent our putative reconsolidation transcriptional program.

**Figure 3.**
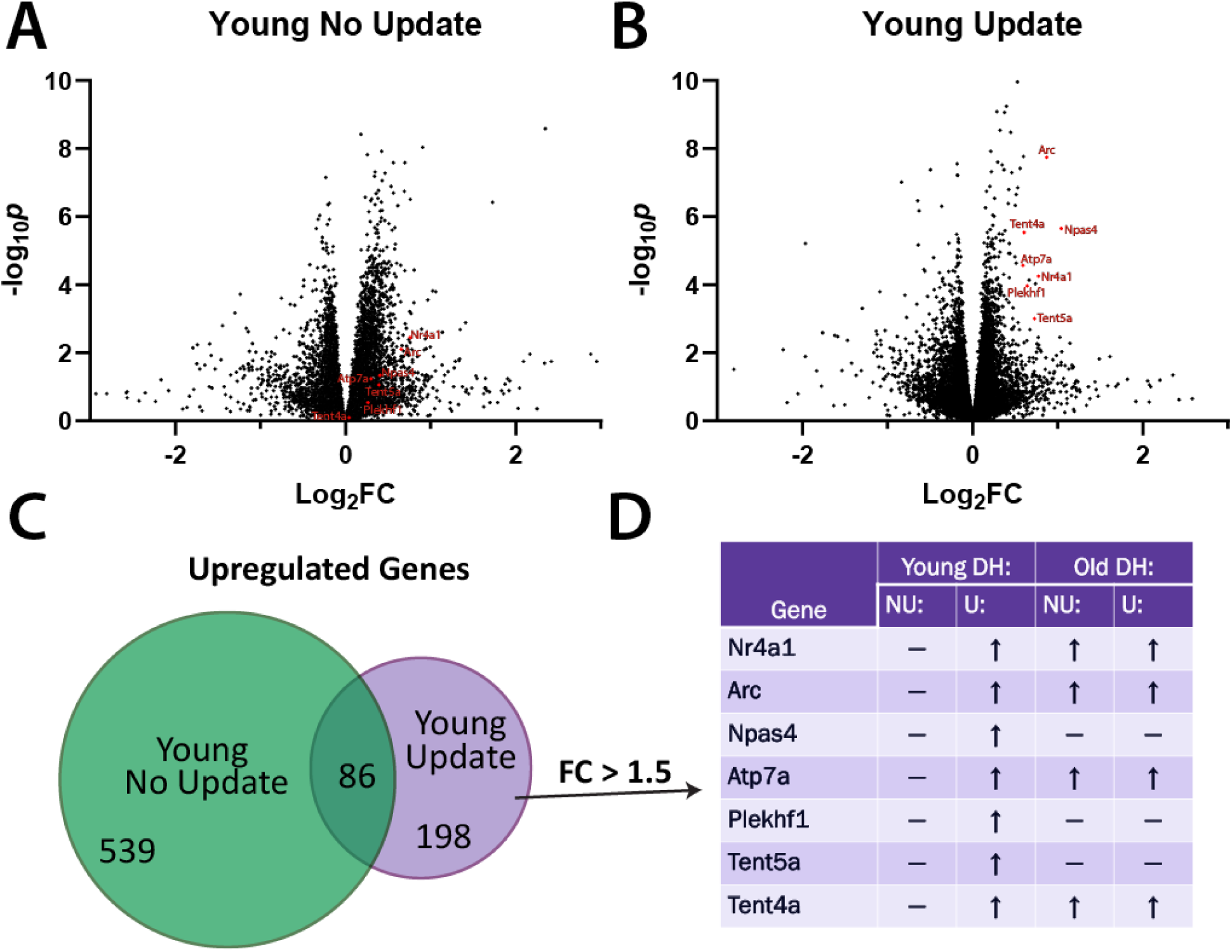
Comparison of DEGs detected in the young NU and U groups. A &. **B.** Volcano plots demonstrating DEGs observed in the young NU and U groups, respectively. Update-specific genes with a fold change > 1.5 are identified in red. **C.** Venn diagram summarizing the overlap in upregulated DEGs detected in the young NU and U groups. **D.** Table summarizing the patterns of expression for the 7 update-specific genes with a fold change > 1.5.

We utilized gene ontology (GO) analyses to examine which biological processes (**Table 1**) and cellular compartments (**Table 2**) were implicated by these lists of DEGs. In the young adult NU group, we observed GO results related to gene silencing, including an enrichment for genes related to miRNA-mediated gene silencing by inhibition of translation (GO:0035278), P-body assembly (GO:0033962), and the RISC complex (GO:0016442). In the young U group, we found multiple GO terms related to Notch signaling, including CSL-Notch-Mastermind transcription factor complex (GO:1990433) and MAML1-RBP-Jkappa-ICN1 complex (GO:0002193), consistent with previous reports of a role for Notch signaling in neuroplasticity and spatial learning (Costa et al., 2003).

**Table 1.** Young biological process GO. Gene ontology biological process results for upregulated DEGs in the young DH, shown separately for the No Update (NU) and Update (U) groups. Results are sorted by enrichment.

| No Update |  |  | Update |  |  |
| --- | --- | --- | --- | --- | --- |
| <u>Biological Process</u> | <u>Enrichment</u> | <u>FDR</u> | <u>Biological Process</u> | <u>Enrichment</u> | <u>FDR</u> |
| external genitalia morphogenesis | 35.22 | 1.05E-02 | positive regulation of dense core granule exocytosis | 76.89 | 1.56E-02 |
| positive regulation of RNA polymerase II transcription preinitiation complex assembly | 15.09 | 2.33E-02 | negative regulation of cytoplasmic translational elongation | 76.89 | 1.55E-02 |
| P-body assembly | 11.01 | 2.28E-02 | receptor-mediated endocytosis involved in cholesterol transport | 51.26 | 3.32E-02 |
| mRNA splice site recognition | 10.44 | 2.23E-02 | positive regulation of potassium ion import across plasma membrane | 23.07 | 1.96E-02 |
| miRNA-mediated gene silencing by inhibition of translation | 10.36 | 1.70E-02 | neuron cell-cell adhesion | 23.07 | 1.95E-02 |

**Table 2.** Young cellular component GO. Gene ontology cellular component results for upregulated DEGs in the young DH, shown separately for the No Update (NU) and Update (U) groups. Results are sorted by enrichment.

| No Update |  |  | Update |  |  |
| --- | --- | --- | --- | --- | --- |
| <u>Cellular Component</u> | <u>Enrichment</u> | <u>FDR</u> | <u>Cellular Component</u> | <u>Enrichment</u> | <u>FDR</u> |
| integrin alphav-beta8<br>complex | 35.22 | 3.00E-02 | extrinsic component<br>of neuronal dense<br>core vesicle<br>membrane | 76.89 | 1.05E-02 |
| interchromatin<br>granule | 21.13 | 9.31E-03 | RAVE complex | 51.26 | 2.33E-02 |
| paraspeckles | 15.65 | 4.01E-03 | CSL-Notch-Mastermind<br>transcription factor<br>complex | 51.26 | 2.28E-02 |
| annulate lamellae | 15.09 | 2.83E-02 | MAML1-RBP-Jkappa- ICN1<br>complex | 51.26 | 2.23E-02 |
| RISC complex | 15.09 | 1.18E-04 | insulin-responsive<br>compartment | 20.97 | 1.70E-02 |

To further narrow down this list of genes, we filtered for DEGs with a fold change greater than 1.5 relative to the HC group, leaving us with a list of 7 update-specific DEGs (**Fig. 3D**; **Table S3**) that exhibited a fold change greater than 1.5 in response to memory updating. Interestingly, each of these 7 genes were either upregulated in neither the old NU or old U group (“aberrantly off” in the old DH during updating: 3 genes) or upregulated in both of these groups (“aberrantly on” in the old DH irrespective of experience: 4 genes), further highlighting the pattern of increased transcriptional similarity between the old U and old NU groups. We focused our investigation on the 3 aberrantly off genes, as they represent the most promising therapeutic targets for targeting age-related cognitive decline by restoring gene or protein expression in the DH.

The 3 genes we identified that were update-specific in the young DH and unchanged in the old brain were *Npas4*, *Plekhf1*, and *Tent5a*. *Npas4* is a known neuron-specific immediate-early gene (IEG; Sun and Lin, 2016) that has repeatedly been shown to play a role in memory formation (Weng et al., 2018; Brito et al., 2024) and has also previously been implicated in memory reconsolidation (Ploski et al., 2011). *Plekhf1* is a gene about which little is known other than that it may help regulate the lysosomal system (Zhu et al., 2023). *Tent5a* (aka *Fam46a* (Warkocki et al., 2018)) encodes for a noncanonical poly(A) polymerase that polyadenylates RNA to improve stability, and is found in both the cytosol and nucleus (Lacidogna et al., 2025). TENT5A is known to stabilize transcripts for secreted proteins in the skeletal (Gewartowska et al., 2021) and immune systems (Liudkovska et al., 2022; Krawczyk et al., 2025), but its role in neuronal function during memory formation or updating is unexplored to date. As proper RNA expression (and therefore, stability) is critical for memory consolidation and reconsolidation, we were interested in further exploring the role of *Tent5a* in memory updating.

### Identification of upregulated genes in the old DH during updating

Thus far, we have investigated reconsolidation-specific genes in the young DH that might underlie memory updating. However, it is also possible that memory-restrictive genes might be aberrantly upregulated during memory updating in the old DH (Abel et al., 1998), thereby inhibiting the learning of the update session. Therefore, we also explored genes that were abnormally upregulated in response to updating in old mice to look for memory repressive genes that might negatively regulate memory updating in the old brain.

We identified 872 upregulated DEGs in the old NU group (**Fig. 4A**; 407 unique to NU) and 818 in the old U group (**Fig. 4B**; 461 unique to U), with 411 DEGs upregulated in both groups (**Fig. 4C**). As with the young mouse DEGs, we used GO analyses to examine which biological processes (**Table 3**) and cellular compartments (**Table 4**) were implicated in these lists of DEGs. In the old DH results, we observed an enrichment of GO terms associated with histone deacetylation, including the Set3 complex (GO:0034967) and the Rpd3L-Expanded complex (GO:0070210) in the old NU group. These results are consistent with a large body of literature linking histone deacetylation to age-related cognitive decline (McQuown and Wood, 2011; Kwapis et al., 2018, 2019b), including a report from our own lab illustrating this process also contributes to age-related memory updating deficits (Smies et al., 2024). Interestingly, we also observed an enrichment for the MOZ/MORF histone acetyltransferase complex (GO:0070776) in the old U group, suggesting that upregulation of histone acetylation-related processes occurs in the old brain during learning, but likely not at the level required to outcompete ongoing histone deacetylation processes.

**Figure 4.**
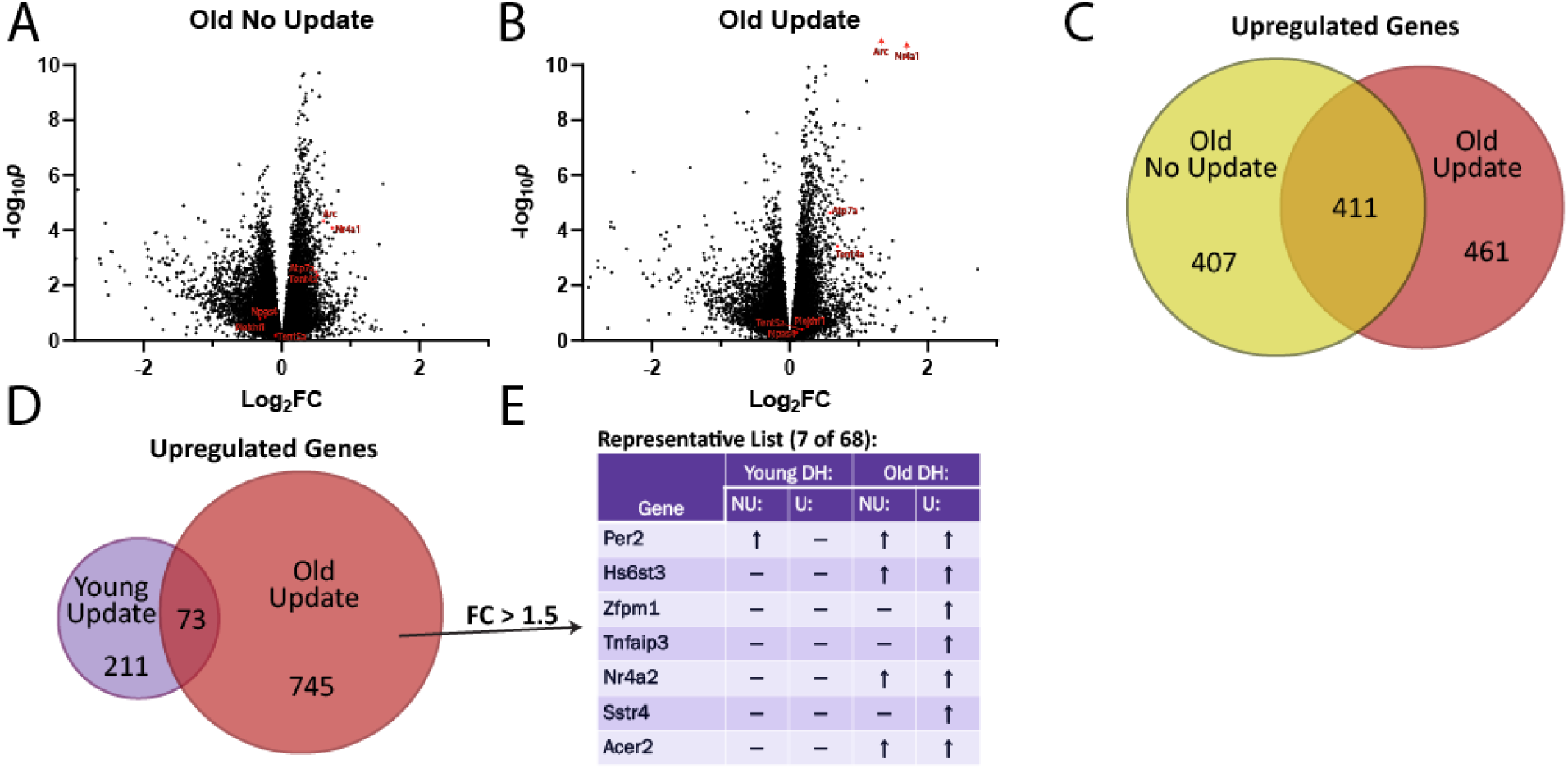
Comparison of DEGs detected in the old NU and U groups. A &. **B.** Volcano plots demonstrating DEGs observed in the old NU and U groups, respectively. Young-mouse update-specific genes with a fold change > 1.5 (from Fig. 3) are again identified in red. **C.** Venn diagram summarizing the overlap in upregulated DEGs detected in the old NU and U groups. **D.** Venn diagram summarizing the overlap in upregulated DEGs detected in the young U and old U groups. **E.** Table summarizing the patterns of expression for a representative list of 7 of the 68 DEGs upregulated in the old U group but not in the young U group exhibiting a large fold change.

**Table 3.** Old biological process GO. Gene ontology biological process results for upregulated DEGs in the old DH, shown separately for the No Update (NU) and Update (U) groups. Results are sorted by enrichment.

| No Update |  |  | Update |  |  |
| --- | --- | --- | --- | --- | --- |
| <u>Biological Process</u> | <u>Enrichment</u> | <u>FDR</u> | <u>Biological Process</u> | <u>Enrichment</u> | <u>FDR</u> |
| heterochromatin<br>boundary formation | 25.21 | 4.21E-02 | cerebellar molecular<br>layer morphogenesis | 26.5 | 3.53E-02 |
| susceptibility to T<br>cell mediated<br>cytotoxicity | 25.21 | 4.20E-02 | retrograde axonal<br>protein transport | 26.5 | 3.53E-02 |
| regulation of<br>synaptic plasticity by<br>chemical substance | 25.21 | 4.20E-02 | positive regulation of<br>branching<br>morphogenesis of a<br>nerve | 26.5 | 3.52E-02 |
| positive regulation of<br>pyruvate<br>decarboxylation to<br>acetyl-CoA | 25.21 | 4.19E-02 | cellular stress<br>response to acid<br>chemical | 26.5 | 3.51E-02 |
| positive regulation of<br>axon extension<br>involved in<br>regeneration | 25.21 | 4.18E-02 | cellular response to<br>Thyroglobulin<br>triiodothyronine | 26.5 | 1.48E-04 |

**Table 4.** Old cellular component GO. . Gene ontology cellular component results for upregulated DEGs in the old DH, shown separately for the No Update (NU) and Update (U) groups. Results are sorted by enrichment.

| No Update |  |  | Update |  |  |
| --- | --- | --- | --- | --- | --- |
| <u>Cellular Component</u> | <u>Enrichment</u> | <u>FDR</u> | <u>Cellular Component</u> | <u>Enrichment</u> | <u>FDR</u> |
| Rpd3L-Expanded complex | 25.21 | 2.87E-02 | AIP1-IRE1 complex | 26.5 | 2.33E-02 |
| Set3 complex | 25.21 | 2.85E-02 | parallel fiber | 26.5 | 2.31E-02 |
| RISC-loading complex | 15.76 | 1.65E-04 | dendritic branch | 18.93 | 5.17E-05 |
| postsynaptic early endosome | 15.13 | 1.25E-02 | MOZ/MORF histone acetyltransferase complex | 15.14 | 1.55E-03 |
| super elongation complex | 15.13 | 1.24E-02 | spectrin-associated cytoskeleton | 15.14 | 1.53E-03 |

We were chiefly interested in identifying genes that were aberrantly upregulated in the old U group compared to their young counterparts, regardless of their behavior in the old NU group. Therefore, we also compared the complete list of all genes upregulated in the old U group (818 total) with the genes upregulated in the young U group (284 total) to identify 73 genes that were upregulated by updating in both age groups, 211 that were upregulated uniquely in the young U group, and 745 that were upregulated uniquely in the old U group (**Fig. 4D**). We then used the same fold change > 1.5 criteria as before to further filter our list of putative updating-repressive genes down to just 68 genes (**Fig. 4E**; **Table S4**). This list contains several interesting candidate DEGs that might be repressing memory updating and follow-up investigations into how these genes affect memory updating are currently ongoing.

Finally, we cross-referenced the lists of DEGs detected across all our comparisons with one another (**Fig. S2**). As our old mice fail to engage a unique transcriptional response to the memory update, we expected both the old U and old NU groups to engage similar genes to the young NU group, reflecting a failure to detect new information and a consistent retrieval-induced transcriptional program. Surprisingly, we noted minimal overlap in either upregulated or downregulated DEGs between the old U group and the young NU group, indicating that age-related memory updating deficits do not simply reflect engagement of the typical NU transcriptional program in response to the update session and instead represent a unique transcriptional response in the old mice.

### Knocking down *Tent5a* in the DH of young mice disrupts memory updating

The most promising candidate gene we identified in our analysis was the noncanonical poly(A) polymerase *Tent5a.* To investigate the role of hippocampal *Tent5a* expression in reconsolidation-mediated memory updating, we used siRNA to knock down *Tent5a* in the DH of young adult mice during the OUL update session. Here, young adult mice underwent the OUL protocol, and 24 hrs after the final training session, an Accell SMARTpool of siRNAs (either a pool targeting *Tent5a* or a non-targeting control pool) was stereotaxically injected into the DH. Forty-eight hrs later, to ensure maximal knockdown (Alaghband et al., 2018), these animals underwent the OUL update session, followed by the test session another 24 hrs later (**Fig. 5A**).

**Figure 5.**
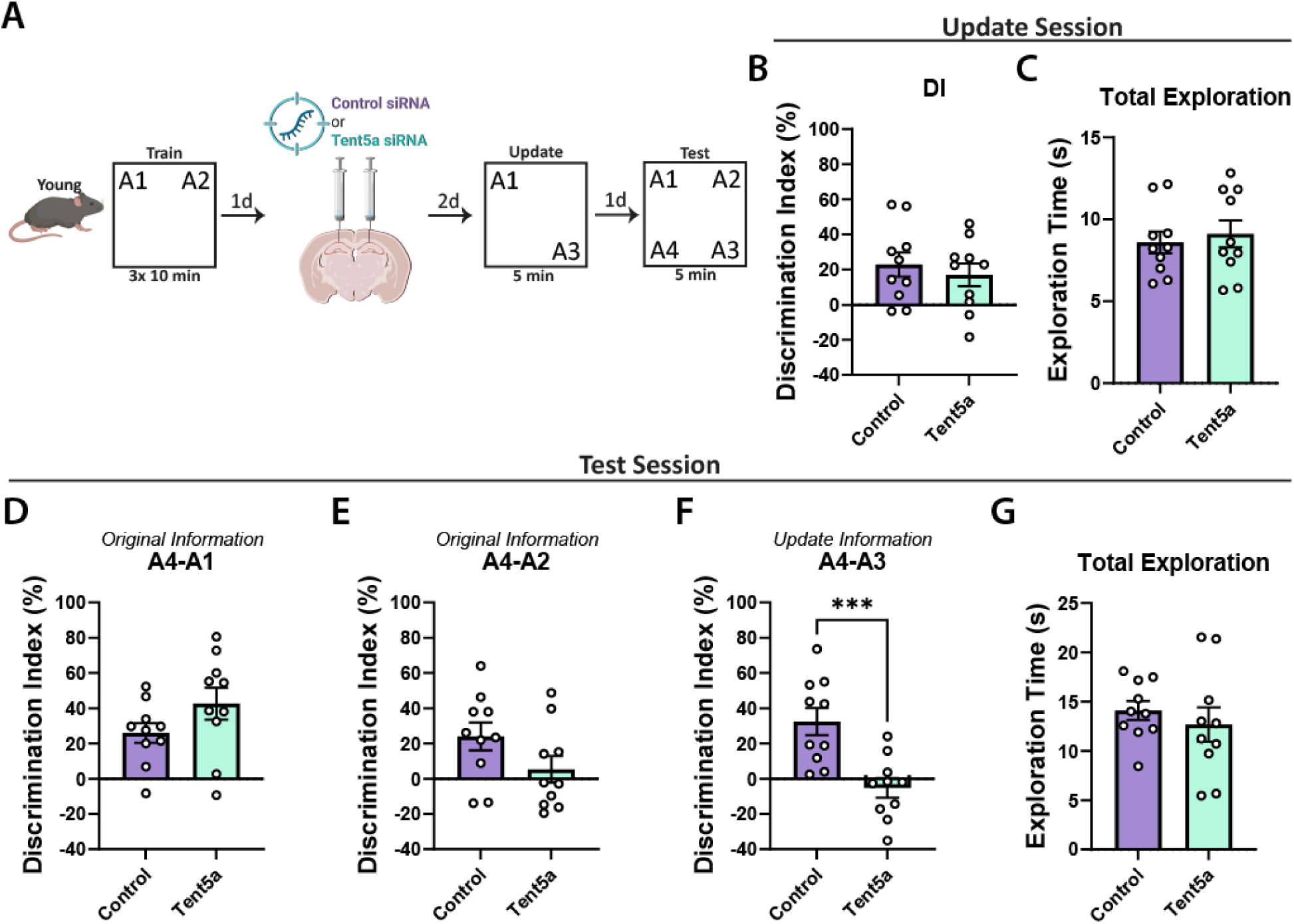
Hippocampal *Tent5a* knockdown disrupts memory updating in young mice. **A.** Experimental schematic. Mice were trained in OUL and then administered either nontargeting control siRNA or siRNA targeting *Tent5a*. Two days later, mice underwent the OUL update and were then tested the following day. **B.** Discrimination indices for training memory assessed during the update session. Both groups of animals showed intact memory for the initial training (one-sample t-tests of A3-A1 DIs against 0: control siRNA: t_9_ = 3.30, p = 0.0093; *Tent5a* siRNA: t_9_ = 2.61, p = 0.028), with no differences between groups (unpaired Student’s t-test: t_18_ = 0.62, p = 0.54). **C.** Total exploration time during the update session. No differences were observed between groups (t_18_ = 0.50, p = 0.62). **D.** During the test session, both groups of mice demonstrated intact memory for training position A1 (one-sample t-tests of A4-A1 DIs against 0: control siRNA: t_9_ = 4.65, p = 0.0012; *Tent5a* siRNA: t_9_ = 4.69, p = 0.0011), with no differences between groups (unpaired Student’s t-test: t_18_ = 1.56, p = 0.14). **E.** Only control mice exhibited intact memory for training position A2 at test (one-sample t-tests of A4-A2 DIs against 0: control siRNA: t_9_ = 3.04, p = 0.014; *Tent5a* siRNA: t_9_ = 0.70, p = 0.50), though a direct comparison detected no differences between groups (unpaired Student’s t-test: t_18_ = 1.72, p = 0.10). **F.** At test, only mice in the control group remembered update position A3 (one-sample t-tests of A4-A3 DIs against 0: control siRNA: t_9_ = 4.19, p = 0.0023; *Tent5a* siRNA: t_9_ = 0.94, p = 0.37), and these mice performed significantly better than mice who received the *Tent5a* knockdown (unpaired Student’s t-test: t_18_ = 3.95, p = 0.0009). **G.** These effects were not driven by any differences in total exploration during the test session (unpaired Student’s t-test: t_18_ = 0.72, p = 0.48). All data are presented as mean ± SEM. n = 10 (4F), 10 (5F).

During habituation, both groups of mice acclimated to the arena as expected (**Fig. S3A**). During the training sessions, mice behaved as expected, with no differences observed in discrimination indices (**Fig. S3B**) or total exploration time (**Fig. S3C**) between groups. During the update session, both groups demonstrated intact memory for the initial training (**Fig. 5B**; one-sample t-tests of A3-A1 DIs against 0: control siRNA: t_9_ = 3.30, p = 0.0093; *Tent5a* siRNA: t_9_ = 2.61, p = 0.028), with no differences between them in terms of DIs (unpaired t-test: t_18_ = 0.62, p = 0.54) or exploration times (**Fig. 5C**; unpaired t-test: t_18_ = 0.50, p = 0.62).

During the test session, both groups of mice demonstrated intact memory for location A1 from training (**Fig. 5D**; one-sample t-tests of A4-A1 Dis against 0: control siRNA: t_9_ = 4.65, p = 0.0012; *Tent5a* siRNA: t_9_ = 4.69, p = 0.0011) with no significant differences between these two groups (unpaired t-test: t_18_ = 1.56, p = 0.14). Control animals also demonstrated intact memory for location A2 from training, while *Tent5a* siRNA animals did not (**Fig. 5E**; one-sample t-tests of A4-A2 DIs against 0: control siRNA: t_9_ = 3.04, p = 0.014; *Tent5a* siRNA: t_9_ = 0.70, p = 0.50) though no significant differences were observed between these groups (unpaired t-test: t_18_ = 1.72, p = 0.10), suggesting *Tent5a* knockdown may have disrupted the memory for location A2. Notably, only the control siRNA group demonstrated intact memory for update location A3 (**Fig. 5F**; one-sample t-tests of A4-A3 Dis against 0: control siRNA: t_9_ = 4.19, p = 0.0023; *Tent5a* siRNA: t_9_ = 0.94, p = 0.37), and there was a significantly higher A4-A3 DI in the control group relative to the *Tent5a* siRNA group (unpaired Student’s t-test: t_18_ = 3.95, p = 0.0009), indicating that hippocampal *Tent5a* knockdown disrupted memory updating in these animals. We observed no differences in total time spent investigating at test (**Fig. 5G**; unpaired Student’s t-test: t_18_ = 0.72, p = 0.48), indicating that these effects were not due to changes in movement at test. Test session exploration time is additionally presented as raw percentages in **Fig. S3D**.

## Discussion

Although the molecular mechanisms underlying memory reconsolidation have been of interest for some time (Tronson and Taylor, 2007; Lattal and Wood, 2013; Bellfy and Kwapis, 2020; Jardine et al., 2022; Brunswick et al., 2025), we still know very little about the transcriptional program required for this process and less still about how this program is affected by aging. Here, we used RNA sequencing combined with the OUL memory updating paradigm to investigate the transcriptional changes that occur following either a memory retrieval session or a reconsolidation-mediated memory update in the young adult and old DH.

In the young adult DH, we expected to observe a large shared transcriptional program between the NU and U groups representing general mechanisms underlying memory retrieval. However, we unexpectedly observed highly distinct transcriptional programs in the NU and U groups (**Fig. 2A** & **3C**), with only a fraction of DEGs (86 out of 823) upregulated by both experiences, and this shared set contained many known IEGs, including *Fos*, *Zif268*, and *Homer1*. Thus, it seems that only a few genes are nonspecifically upregulated in response to memory retrieval, but a large set of genes is engaged specifically by reconsolidation (198 total genes), and an even larger set is upregulated when a memory is retrieved without initiating reconsolidation (539 total genes). This suggests that memory retrieval without updating is an active cellular process in which the DH engages an expansive list of genes, possibly to prevent the memory destabilization and synaptic remodeling that occurs during memory reconsolidation (see Tronson and Taylor, 2007; Bellfy and Kwapis, 2020; Jardine et al., 2022). This position is consistent with the enrichment of GO terms associated with gene silencing in this group (**Tables 1**, **2**), suggesting that active silencing of reconsolidation-related genes may occur in the absence of prediction error.

In the old DH, we observed a greater overlap (411 genes) between upregulated DEGs in the NU (872 total genes) and U groups (818 total genes), indicating these two experiences engage similar transcriptional programs. We also observed greater similarities in old NU and old U log_2_ fold change for specific genes in the old DH (R^2^ = 0.51, slope = 0.69) than in the young DH (R^2^ = 0.13, slope = 0.35), further indicating an age-related increase in transcriptional similarity. Additionally, none of the 7 putative update-specific DEGs identified in the young brain exhibited a behavior-specific change in the old brain; each gene was either upregulated in neither the old NU nor old U group or upregulated in both groups. Although this might indicate that the old DH, unlike the young DH, operates a single shared retrieval transcriptional program common to both groups, the more likely explanation is that this increase in transcriptional similarity is contributing to age-related updating deficits, reflective of the known increases in pattern completion occurring in the old brain (Yassa and Stark, 2011; Cès et al., 2018), which might extend even to the level of individual transcripts. Neither the old NU group nor the old U group demonstrate intact memory for the update session the following day (Kwapis et al., 2019a), though old mice are capable of acquiring the update information into short-term storage (Brunswick et al., 2026), making the transcriptional similarities reported here in line with previously observed behavioral similarities.

The presence of several IEGs in these gene lists is unsurprising, as these genes have long been known to be engaged in response to both memory formation and retrieval. *Zif268* has previously been suggested to play an important role in memory reconsolidation (Lee et al., 2004), suggesting it might appear uniquely in our young U group. The fact that we observed upregulation of this gene not only in the young U but also in the young NU group (which does not undergo reconsolidation (Kwapis et al., 2019a)), suggests that *Zif268* may be related to memory retrieval more broadly rather than specifically necessary for reconsolidation, consistent with other work indicating a more general role of *Zif268* in memory processes (e.g., Kwapis et al., 2018; Trask et al., 2020). The identification of the IEG *Npas4* uniquely in the young U group is in line with prior work demonstrating that this gene contributes to learning broadly (Sun and Lin, 2016; Weng et al., 2018; Brito et al., 2024) and may specifically play a role in memory reconsolidation (Ploski et al., 2011). Prior work has also shown that *Npas4* is both important for maintaining excitatory-inhibitory balance by regulating incoming inhibition (Lin et al., 2008) and responsible for promoting memory discrimination (that is, pattern separation) within the DH (Sun et al., 2020). Together, this evidence suggests that high *Npas4* expression during reconsolidation may remodel the training engram to incorporate the update information, perhaps by increasing inhibition onto some of the training-associated engram neurons. Along the same lines, the lack of *Npas4* induction in the old U group might contribute to the decline in pattern separation typically observed in age. We demonstrated for the first time a role for *Tent5a* in learning and memory. Although *Tent5a* has previously been linked to cognition in two genome-wide association studies (Nicodemus et al., 2014; Sommerer et al., 2023), this is the first study to investigate a causal role of this gene in learning and memory, specifically showing that DH expression of *Tent5a* is necessary for memory reconsolidation. *Tent5a* encodes a noncanonical poly(A) polymerase that adds adenosine tails to mature mRNA (Lacidogna et al., 2025). As both transcription and translation are necessary for memory reconsolidation, one possibility is that TENT5A polyadenylates other reconsolidation-associated mRNAs to promote stability and increase the likelihood that these transcripts are translated during reconsolidation. In our study, we did not examine the subcellular localization of either *Tent5a* transcripts or TENT5A protein during reconsolidation. Perhaps TENT5A—which is found in the cytoplasm unlike most canonical poly(A) polymerases (Krawczyk et al., 2025)—specifically plays a role in maintaining the stability of transcripts bound for local translation at the synapse. TENT5A also plays an important role in regulating secreted proteins (Lacidogna et al., 2025), and, in neurons, it might control the secretion of either neuropeptides or components of the extracellular matrix, both of which play critical roles in learning and memory. It reasons that TENT5A could therefore play a critical role in orchestrating the stability and translation of key transcripts important for reconsolidation-dependent memory updating.

Although our work here shows that *Tent5a* is necessary for successful memory reconsolidation, we have not yet tested whether *Tent5a* is also necessary for consolidation of the initial memory. Interestingly, *Tent5a* was implicated in a previous RNA sequencing study from our lab investigating the transcriptional mechanisms necessary for initial memory consolidation in young mice specifically during the daytime, when spatial memory is best (Bellfy et al., 2023). Thus, we would expect *Tent5a* to play an important role in memory consolidation, as well.

We are also interested in further exploring the putative update repressive genes identified in this study, many of which are well-poised to play an important role in memory processes. For instance, in our old U group we detected an upregulation of genes central to the circadian clock, which has a well-established role in mediating cognitive performance (Kwapis et al., 2018; Smies et al., 2022; Brunswick et al., 2023); genes responsible for modifying heparan sulfate, which has been implicated in the development of Alzheimer’s disease (Snow et al., 1988; Schultheis et al., 2024); and genes broadly linked to inflammation, which is a well-established mediator of age-related cognitive decline (Mekhora et al., 2024). We found, for instance, that the old U group exhibited upregulation of *Tnfaip3* (tumor necrosis factor alpha-induced protein 3), a known repressor of NF-κB, which is a transcription factor known to play an important role in learning and memory (Crampton and O’Keeffe, 2013; Kaltschmidt and Kaltschmidt, 2015). Interestingly, *Tnfaip3* has also been shown to activate microglia (Voet et al., 2018), which play important roles in synaptic plasticity (Sipe et al., 2016), learning (Parkhurst et al., 2013; Nguyen et al., 2020; Liu et al., 2026), and forgetting (Wang et al., 2020). Furthermore, microglia become sensitized with age, entering into a pro-inflammatory state that interrupts their ability to properly interact with nearby neurons (Norden and Godbout, 2013). Thus, it seems that *Tnfaip3* upregulation could be disrupting memory reconsolidation in the aged DH by inhibiting NF-κB within engram neurons and/or excessively activating microglia and thereby disrupting synaptic remodeling occurring during memory reconsolidation. Future experiments will look to see which cell types in the old brain induce expression *Tnfaip3* in response to memory updating and if hippocampal *Tnfaip3* knockdown is sufficient to improve memory updating in old mice.

In sum, these results explore age-related changes in hippocampal transcription during memory updating, both revealing broad changes in transcriptional patterns that occur during updating and identifying specific genes (for instance, *Tent5a*) that seem to play important but previously unexplored roles in memory reconsolidation. Future experiments will expand on these results to identify other memory-relevant genes (e.g., potentially *Tnfaip3*) while also further elucidating the mechanism by which *Tent5a* mediates memory updating.

---

**Supplemental Figure 1.**
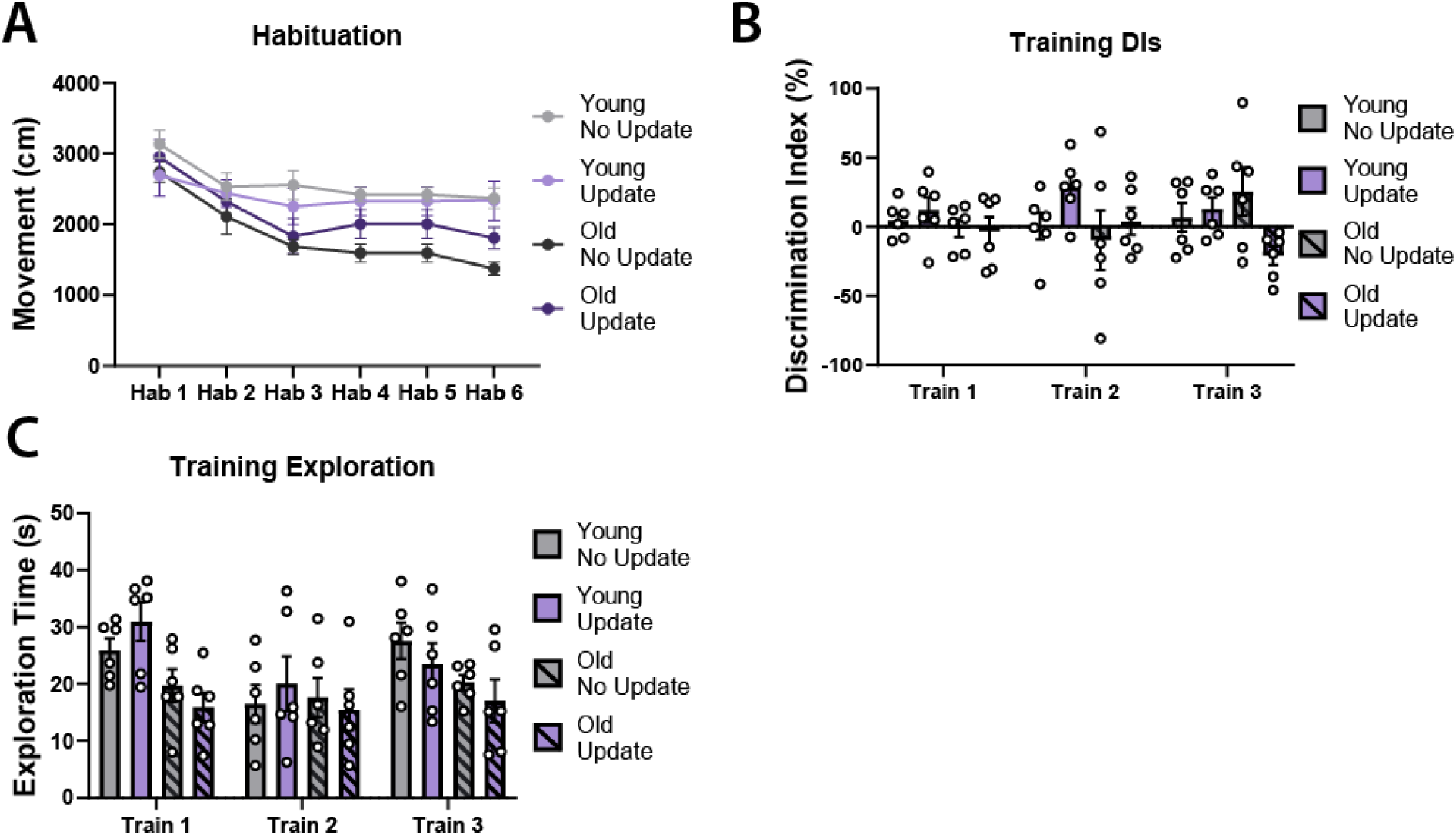
**Additional behavioral data from the RNA Sequencing experiment (Figure 1). A.** All groups habituated to the arenas as expected, with decreased movement observed in old mice (mixed-effect analysis: effect of session: F_(2.94,58.25)_ = 28.99, p < 0.0001; effect of age: F_(1,20)_ = 9.07, p = 0.0069; effect of behavior: F_(1,20)_ = 0.19, p = 0.67; session x age interaction: F_(2.94,58.25)_ = 4.25, p = 0.0092; session x behavior interaction: F_(2.94,58.25)_ = 1.047, p = 0.38; age x behavior interaction: F_(1,20)_ = 2.39, p = 0.14; session x age x behavior interaction: F_(2.94,28.25)_ = 0.20, p = 0.90). **B.** No group exhibited a significant object preference during the training sessions, though we did detect a session x behavior interaction (three-way RM-ANOVA: effect of session: F_(1.85, 37.08)_ = 0.083, p = 0.91; effect of age: F_(1, 20)_ = 3.20, p = 0.089; effect of behavior: F_(1, 20)_ = 0.025, p = 0.88; session x age interaction: F_(1.85, 37.08)_ = 0.21, p = 0.80; session x behavior interaction: F_(1.85, 37.08)_ = 3.39, p = 0.048; age x behavior interaction: F_(1, 20)_ = 3.73, p = 0.068; session x age x behavior interaction: F_(1.85, 37.08)_ = 1.099, p = 0.34). **C.** Both groups spent less time investigating across training sessions, with old mice exploring less overall (three-way RM-ANOVA: effect of session: F_(1.62, 32.42)_ = 3.75, p = 0.043; effect of age: F_(1, 20)_ = 10.08, p = 0.0048; effect of behavior: F_(1, 20)_ = 0.15, p = 0.71; session x age interaction: F_(1.62, 32.42)_ = 2.032, p = 0.15; session x behavior interaction: F_(1.62, 32.42)_ = 0.6233, p = 0.51; age x behavior interaction: F_(1, 20)_ = 1.267, p = 0.27; session x age x behavior interaction: F_(1.62, 32.42)_ = 0.6204, p = 0.51).

**Supplemental Figure 2.**
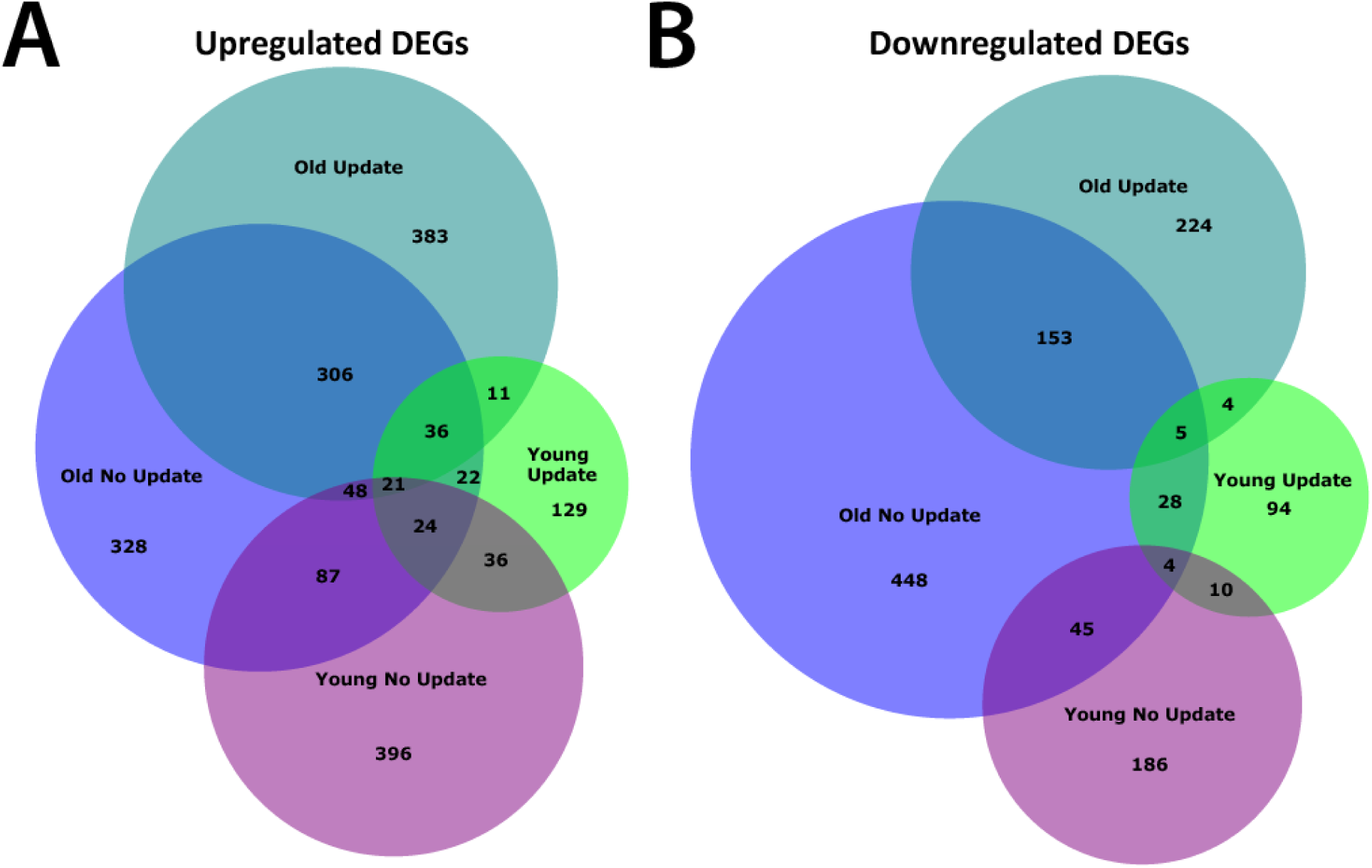
Patterns of expression for all DEGs across age groups. **A.** Venn diagram summarizing all upregulated DEGs detected across all 4 groups and the overlaps between them. **B.** Venn diagram summarizing all downregulated DEGs detected across all 4 groups and the overlaps between them.

**Supplemental Figure 3.**
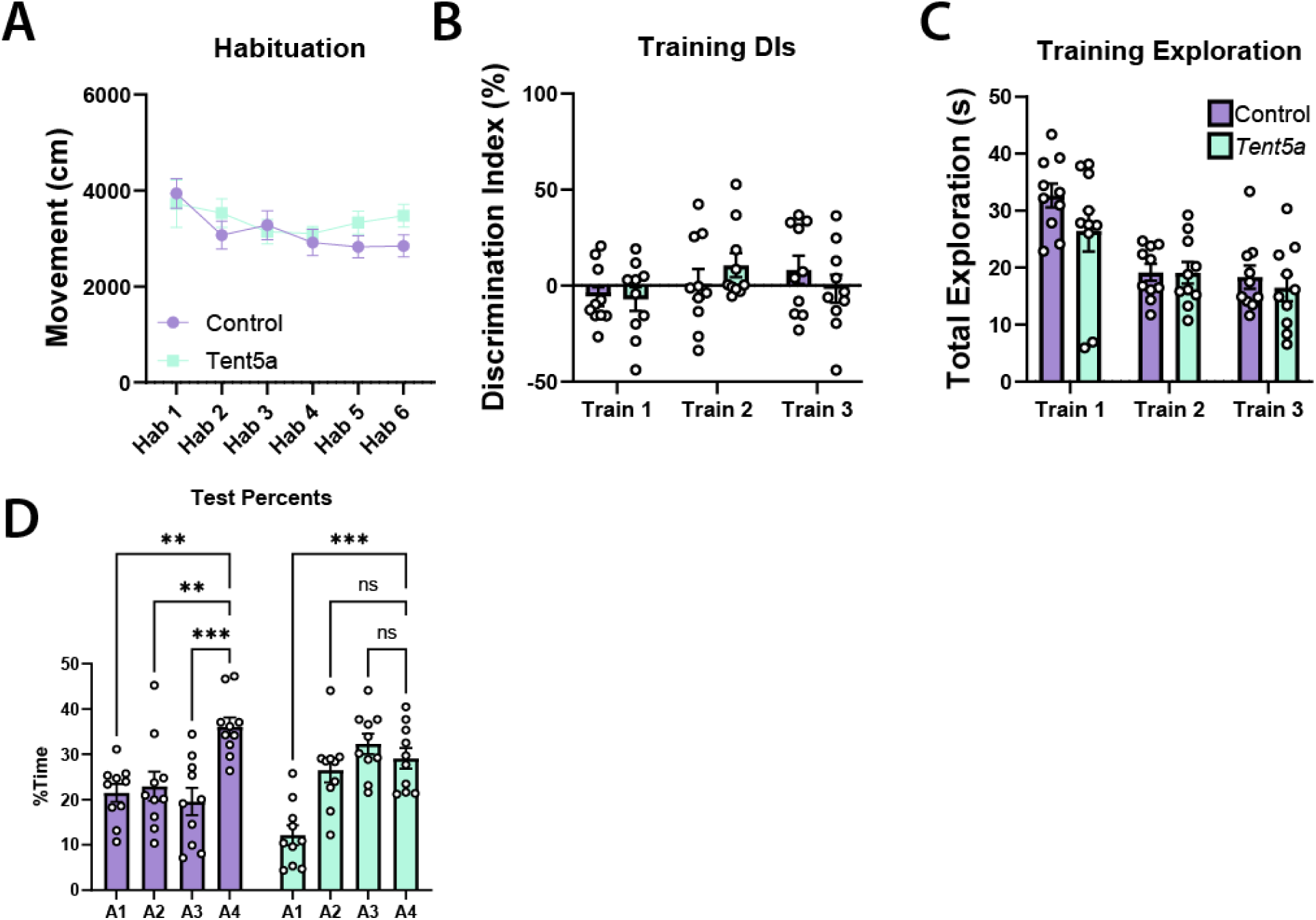
**Additional behavioral data from the Tent5a siRNA experiment (Figure 5). A.** Both groups habituated to the arenas as expected (two-way RM ANOVA: effect of session: F_(5,90)_ = 4.81, p = 0.0006; effect of siRNA: F_(1,18)_ = 0.55, p = 0.47; effect of subject: F_(18,90)_ = 8.60, p < 0.0001; interaction: F_(5,90)_ = 1.71, p = 0.14). **B.** Neither group had a significant object preference during the training sessions (two-way RM ANOVA: effect of session: F_(2,36)_ = 2.69, p = 0.081; effect of siRNA: F_(1,18)_ = 0.0055, p = 0.94; effect of subject: F_(18,36)_ = 2.30, p = 0.017; interaction: F_(2,36)_ = 1.56, p = 0.22). **C.** Both groups spent comparable time investigating objects during training (two-way RM ANOVA: effect of session: F_(2,36)_ = 21.91, p < 0.0001; effect of siRNA: F_(1,18)_ = 1.31, p = 0.27; effect of subject: F_(18,36)_ = 2.19, p = 0.022; interaction: F_(2,36)_ = 1.27, p = 0.29). **D.** Investigation times at test displayed as percentages two-way RM ANOVA: effect of object: F_(3,54)_ = 9.98, p < 0.0001; effect of siRNA: F_(1,18)_ = 1.68, p = 0.21; effect of subject: F_(18,54)_ = 3.77 x 10^-14^, p > 0.999; interaction: F_(3,54)_ = 6.19, p = 0.0011; * denotes p < 0.05, ** denotes p < 0.01, *** denotes p < 0.001, **** denotes p < 0.0001 on Dunnett’s multiple comparisons test against A4). All data are presented as mean ± SEM.

Supplemental Table 1.

**All young DEGs.** List of all differentially expressed genes (upregulated or downregulated) in the young mouse DH in response to the no update or update session.

Supplemental Table 2.

**All old DEGs**. List of all differentially expressed genes (upregulated or downregulated) in the old mouse DH in response to the no update or update session.

Supplemental Table 3.

**DEGs upregulated in young only by updating**. List of 7 update-specific genes in the young mouse DH exhibiting a large fold change (>1.5).

Supplemental Table 4.

**DEGs upregulated by updating only in old**. List of 68 genes upregulated by updating in the old but not young DH and exhibiting a large fold change (>1.5).

## Author Contributions

C.A.B., J.L.K., and I.A. designed research C.A.B., A.G.D., T.A.W., A.R.M., S.M., P.-H.C., S.M.J., and G.C.P. performed research C.A.B., A.G.D., and A.S., analyzed data. C.A.B. and J.L.K. wrote and edited the paper.

## Conflicts of Interest

No competing financial interests.

## Acknowledgements

This work was supported by National Institutes of Health grants F31AG087533 (C.A.B.), R21AG068444 (J.L.K.), and R01AG074041 (J.L.K.); by the McKnight Brain Research Foundation/AFAR Innovator Award in Cognitive Aging and Memory Loss (J.L.K.); by the Hevolution/AFAR New Investigator Award in Aging Biology and Geroscience Research (J.L.K.) and by the J. Lloyd and Dorothy Foehr Huck Institutes of the Life Sciences at Penn State (A.S. and I.A.). We wish to thank Dr. Craig Praul and the Penn State Huck Genomics Core Facility (RRID:SCR_023645) for conducting all sequencing for the transcriptomics experiment. We would also like to thank all members of the Kwapis Lab for scientific discussion and technical assistance. Figures were made with BioRender. Venn diagrams were made with BioVenn (Hulsen et al., 2008) and DeepVenn (Hulsen, 2022).

## Notes

### Competing Interest Statement

The authors have declared no competing interest.

