## Supplemental Table 1 for "Aging disrupts transcriptional programs for memory updating in the dorsal hippocampus and reveals a required role for *Tent5a* in reconsolidation"

| No Update | | Update | |
|---|---|---|---|
| Down | Up | Down | Up |
| Gm7488 | Ctnnb1 | Srp54b | Bace1 |
| Gm49510 | Cdv3 | Gm8797 | Ttc3 |
| Oasl2 | Abcd2 | Gm10599 | Astn1 |
| Rsc1a1 | Pdcd6ip | Rps27a-ps: | Kif5c |
| Ttc34 | Sec22c | Mgp | Fbxo28 |
| Sebox | Brox | Nhlh1 | D5Ertd579e |
| Tuba8 | Usp8 | Gm17081 | Syt4 |
| Gprc5c | Slc2a3 | Pcdhgc4 | Cdk5r1 |
| Sst | Ddx17 | Efna1 | Nrxn1 |
| Calhm5 | Nalcn | Maskbp3 | Pafah1b1 |
| Snurf | Pak2 | Gm54215 | Atp2c1 |
| Pnoc | Cbx5 | Vmn2r97 | Hif1an |
| Nudt17 | Ythdc1 | Tnfsfm13 | Socs7 |
| Glyctk | Pik3r3 | Bgn | Camsap1 |
| Isoc2b | Cadm1 | Gprc5c | Ddi2 |
| Klk8 | Stxbp4 | Klhl40 | Fbxw11 |
| Cort | Mtss1 | Prkn | Jak1 |
| Cd248 | Dennd4a | Ubxn11 | Tspyl4 |
| Nbl1 | Tnrc6a | Cd37 | Stxbp5 |
| Gm15387 | Med12l | Pear1 | Itch |
| Nhp2 | Prkacb | Fxn | Slc6a8 |
| Selenow | Kdm3a | Myl9 | Gm49336 |
| Pcdhgc4 | Sc5d | Ccdc189 | Jmy |
| Fdx2 | Wdr37 | Col22a1 | Pdcd6ip |
| Dact2 | Ppp1r15b | Krt77 | Irgq |
| Drap1 | Miga1 | Cldn5 | Otud7a |
| Sp2 | Lrig2 | Tyrobp | Ythdf3 |
| Lsm7 | Hnrnpd | Igfbp6 | Acbd3 |
| Aprt | Prkaa1 | Cebpd | Rapgef2 |
| Gamt | Zfp644 | Gm29216 | Acsl3 |
| Zfp414 | Enah | Stxbp2 | Slc36a4 |
| Pwwp2b | Cstf3 | Or2y1 | Ank2 |
| 3110082I1 | Nup205 | Prdm5 | Atrnl1 |
| Rpl41 | Zmpste24 | Gm42742 | Larp4b |
| Saysd1 | 8030462N17Rik | Gm14418 | Klhl3 |
| Msrb1 | Cldn12 | 3110082I1 | Ahdc1 |
| Rpp21 | Cyp51 | Lrrc45 | Mfsd14a |
| Cpxm1 | Fbxo28 | Thbd | Stk4 |
| 1810009A1 | Tgoln1 | Tle2 | Atp8b2 |
| Thoc6 | Sgms1 | Chrd | Abl2 |
| Mcts2 | Dnajc3 | Atosa | Nin |
| Flywch2 | Usp13 | Wsb1 | Cep170 |

| | | | |
|---|---|---|---|
| Mri1 | Sfpq | Penk | Akt3 |
| Aopep | Tardbp | Zfp950 | Wdr7 |
| Lynx1 | Slc1a2 | Ly6a | Nek1 |
| Pet100 | Dock7 | Zcchc7 | Nup153 |
| Sirt6 | Zswim5 | Spata33 | Dnajb1 |
| Aldh1b1 | Kbtbd2 | Bex1 | Atp8a1 |
| Fth1 | Cand1 | Rps19 | Usp32 |
| Mlf2 | Usp37 | Rplp1 | Zcchc2 |
| Ubl7 | Ahctf1 | Rpl31 | Plaa |
| Otub1 | Msmo1 | Gm15500 | Usp31 |
| Thy1 | Tgs1 | Rpsa | Stk24 |
| Cyb5r3 | Tmem41b | Pfdn5 | Fam91a1 |
| Bckdk | Dcbld2 | Rpl7a | Dcp2 |
| Iffo1 | Irf2bp2 | Rps14 | Arfgef1 |
| Npdc1 | Dcp1a | Fau | Scaper |
| Rab1b | Cep97 | Rps25 | Cpeb3 |
| Fkbp8 | Kdm5a | Rps24 | Daam1 |
| Pebp1 | Smarca5 | Rps11 | Ankrd50 |
| Map3k11 | Tjp1 | Rps18 | Psd3 |
| Ubb | Lrba | Rpl19 | Pcnx |
| Syt3 | Atad2b | Rpl41 | Plcl1 |
| Hmox2 | Usp24 | Ramp1 | Slc4a10 |
| Cby1 | A930017M01Rik | Rps27 | Pcdhac2 |
| Naa38 | Pnpla8 | Rpl38 | Gpr158 |
| Zfp346 | Phlpp2 | Sh3bgrl3 | Ubxn7 |
| Dapk3 | Rab21 | Rps28 | Tgoln1 |
| Slc35b2 | Ankrd6 | Ppp1r1a | Togaram1 |
| Scn1b | U2surp | Rps29 | Lrrn3 |
| Ppp1r15a | Ankrd26 | Mrpl40 | Cntn1 |
| Lsm4 | Mkln1 | Rps15 | Slc38a1 |
| Adap1 | Copb1 | Rpl17-ps3 | Lemd3 |
| Dnajc30 | Tial1 | Zfp414 | Usp37 |
| Ndufa5 | Myo6 | Gfra4 | Rlf |
| Cops7a | Ddx6 | Rps5 | E2f3 |
| Rab4b | Atg2b | Rpl23a | Irf2bpl |
| Nat14 | Kif5b | Vps25 | Kcna2 |
| Rae1 | Alkbh5 | 0610010K1 | Luzp1 |
| Rnasek | Ankrd45 | Zc3hc1 | Gtf3c3 |
| Snrpa | Top1 | Gng10 | Mtmr4 |
| Ppp2r1a | Pkd1 | Rpl28 | Prickle1 |
| Inpp5k | Sos2 | Ccl27a | Ubxn2a |
| Timm13 | Paqr8 | H2-Ke6 | Arfgef2 |
| Pex16 | Ssr1 | Tab1 | Stam |
| Uqcr11 | Morc3 | Dbp | Fbxo42 |

| | | | |
|---|---|---|---|
| Ptms | Paxbp1 | Ranbp1 | Ubr3 |
| Pdk2 | Slc35e2 | Rpl37 | Rbpj |
| Ankrd13b | Msi2 | Rps7-ps3 | Cerk |
| Arrb2 | Csmd1 | Ift20 | Camsap2 |
| Wdr24 | Zcchc2 | Ndufa3 | Faxc |
| Gapdh | Usp38 | Gps2 | Cmip |
| Hras | Prkar2b | Cdk10 | Pcdh1 |
| Rnf126 | Prpf4b | Arrb2 | Lrp6 |
| Nabp2 | Zfp780b | Sp2 | Cpeb2 |
| Atp6v1g2 | Smc5 | N4bp2l1 | Gspt1 |
| Rnf167 | Nop58 | Oscp1 | Lrp12 |
| Apbb1 | Isoc1 | Snrnp27 | Myo5a |
| Snf8 | Hdgfl3 | Igfbp7 | Cpd |
| Cstpp1 | Zranb2 | Bnip1 | Cdh8 |
| Taldo1 | Stox2 | Mus81 | Bcor |
| Ap1m1 | Pdgfra | Nrgn | Arid4b |
| Manbal | A830018L16Rik | Ndufb7 | Mcl1 |
| Slc7a4 | Ago4 | Cdk5rap3 | Uhmk1 |
| Gatb | Spag9 | Usp2 | Ankib1 |
| Tbrg1 | Ern1 | Zfp821 | Man1a2 |
| Sh3bgrl3 | Hspa5 | Use1 | Dennd4a |
| Vps25 | Lrp6 | Rnf166 | Epm2aip1 |
| Matk | Sema6d | Crip2 | Hipk3 |
| Ift52 | Setx | Alpk2 | Dock4 |
| Them6 | Smndc1 | Rpl17 | Dmxl2 |
| Ap1s1 | Vav3 | Pex16 | Trip12 |
| Psmb3 | Dnmt3a | Commd4 | Ppm1l |
| Srrd | Esyt2 | Mak16 | Mtmr9 |
| Capzb | Appbp2 | Hdac7 | Larp4 |
| Znhit2 | Gtf3c4 | Rps3a1 | Zswim5 |
| Tcta | Mat2a | Rpl13a | 1810013L24Rik |
| BC005624 | Tulp4 | Rpl14 | Kif1b |
| Lrrfip1 | D430041D05Rik | Rps8 | Zfp719 |
| Hagh | Osbpl11 | Cox5b | Snrk |
| Snx15 | Pik3ca | Rps23 | Rnf165 |
| Smyd5 | Azin1 | Rps16 | Mfap3 |
| 2310033P0 | Agl | Rpl13 | Maml1 |
| Map7d1 | Rnf144b | Rpl8 | Scyl2 |
| Carmil2 | Rmnd5a | Rpl30 | Pom121 |
| Cops9 | Zfp280b | Rps13 | Synj1 |
| Tmem59l | Ptpn12 | Rps4x | Lrrc8a |
| Dctn2 | Mtmr4 | Ndufa5 | Dennd11 |
| Tpst2 | Sdf2l1 | Rps10 | Cdh11 |
| Ruvbl1 | Vamp4 | Rpl24 | Clock |

| | | | |
|---|---|---|---|
| Tomm6 | Ppig | Txn2 | Zfp655 |
| Cfl1 | Usp15 | Rpl21 | Hycc2 |
| Mt3 | Papola | Rpl34 | Sema6d |
| Rpl18a | Bltp1 | Rps2 | Usp9x |
| Dmwd | Slc4a4 | Rpl27 | Srgap3 |
| Gpatch1 | Nup155 | Rpl32 | Prdm2 |
| Rnf187 | Stag1 | Rpl26 | Ssh2 |
| Dnal4 | Khdc4 | Rpl3 | Tnpo1 |
| Cuedc2 | Mtx3 | Rpl37a | Rfx7 |
| Thyn1 | L3mbtl3 | Mpc1 | Zfp280b |
| Fabp3 | Gna13 | Rps6 | Pag1 |
| Chchd6 | Wnk3 | Rpl4 | Elovl6 |
| AI837181 | Dhx15 | Rpl27a | Ubfd1 |
| Babam2 | Pag1 | Sephs2 | Tet3 |
| Mrpl38 | Gm49336 | H3f3b | Wnk3 |
| Kcnab2 | Ankrd17 | Clk3 | Epha5 |
| Bmerb1 | Dync2h1 | Rpl10 | Frmd6 |
| Nosip | Plxdc2 | Rpl36a | Atp13a3 |
| Mcrs1 | Otud6b | Rpl11 | Rb1cc1 |
| Atp6v1e1 | Ubxn2a | | Sacs |
| Katnb1 | Zfp955b | | Pcdhgc5 |
| Aldoa | Irs1 | | Egr3 |
| Park7 | Ddi2 | | Tmem68 |
| Arpc3 | Picalm | | Mtf1 |
| Mrpl52 | Ddx19b | | Ipo7 |
| Thop1 | Agpat5 | | Zbtb33 |
| Zfp821 | Ermp1 | | Mib1 |
| Mcat | Smarca1 | | Cdk7 |
| Mtx1 | Lrrn3 | | Unc5c |
| Denr | Hnrnpll | | Arid2 |
| Rundc3a | Nek1 | | Topbp1 |
| Adck1 | Arglu1 | | Atg2b |
| Mus81 | Ggps1 | | Dock11 |
| Zfp580 | Eea1 | | D130043K22Rik |
| Sdhaf1 | Cep290 | | Mapk8 |
| Gstp1 | Ypel2 | | Adnp2 |
| Cabp1 | Ppp4r2 | | Map4k4 |
| Dus1l | Agps | | Spring1 |
| Ntpcr | Fubp1 | | Wnk1 |
| Mydgf | Mid2 | | Setd7 |
| Sra1 | Ptprz1 | | Zfp516 |
| Cables2 | Slc30a7 | | Vcpip1 |
| Rpl27 | Dcp2 | | Zfp407 |
| Ndufa10 | Slitrk2 | | Ralgapa1 |

| | | |
|---|---|---|
| Msra | Sesn3 | Ranbp2 |
| Scamp4 | Znrf2 | Adipor2 |
| Rps14 | Nup153 | Nalcn |
| Cbr1 | Map2 | Sacm1l |
| Cyth2 | Peg3 | Ankrd17 |
| Chmp6 | Smg1 | Htr2a |
| Vps9d1 | Edem3 | Irs2 |
| Gm10275 | Fmnl2 | Zbtb11 |
| Get3 | Pdzd8 | Hmgcr |
| Gba | Apaf1 | Midn |
| Ankrd34a | Ttbk2 | Apc |
| Med8 | Gmps | Kcna1 |
| Zfp653 | Zfp329 | Atp2a2 |
| Zbtb22 | Hmgcr | Pcdh7 |
| Ubald1 | Irs2 | Bsn |
| Txnrd2 | Rictor | Hdac4 |
| Selenom | Pals1 | Arl5b |
| Tpgs1 | Bmpr1a | Map3k13 |
| Cox6a1 | Lifr | Nufip2 |
| Btbd2 | Ppm1d | Pcf11 |
| Map2k2 | ENSMUSG00000121477 | Nav3 |
| Dynlrb1 | Kcnn3 | Scn2a |
| Ptprn | Cnot6 | Ankrd52 |
| Ruvbl2 | Fam91a1 | Mia3 |
| Nagk | Mcl1 | Lnpep |
| Lrrc20 | Mtmr10 | Wdfy3 |
| Mrps33 | Hif1a | Ldlr |
| Miip | Csmd3 | Brwd3 |
| Idh3g | Dennd4c | Clmn |
| Snrnp27 | Cep350 | Adamts3 |
| Ndufa7 | Zfp638 | Dusp4 |
| Gsto1 | AI987944 | Ranbp6 |
| Ccdc85b | Chl1 | Atm |
| Supt4a | Slc25a32 | Arl4d |
| Tmub1 | Ago3 | Csrnp1 |
| Snrpn | Slc44a5 | Ccdc117 |
| Top1mt | Btbd7 | Card6 |
| Eefsec | Trpm7 | Epha3 |
| Tmem160 | Hook3 | Cep350 |
| Dhrs7 | P4ha1 | Klf12 |
| Tmem234 | Thsd7a | Cadps |
| Btbd6 | Prex2 | Tmcc3 |
| Pld3 | N4bp2l2 | Slc9a7 |
| Ddx49 | Btaf1 | Gm28036 |

| | | |
|---|---|---|
| Aplp1 | Zdhhc21 | Spen |
| Poldip2 | Fndc3b | Edil3 |
| Phactr3 | Zfp292 | Srp54c |
| Ppme1 | Ppp4r3a | Per1 |
| Abhd12 | Patl1 | Sdk2 |
| Eri3 | Slc30a1 | Bach2 |
| Mtch1 | Dmd | Zfp770 |
| Ppp6r2 | Lgr4 | Gm4202 |
| Vamp2 | Socs6 | Dmxl1 |
| Atp5g1 | Arl13b | Ahr |
| Wbp11 | Nab1 | Robo1 |
| Smarcb1 | Lats1 | Rbm12b1 |
| Higd2a | Xpr1 | Slc5a3 |
| Rab7 | Lrrc58 | Pigg |
| Ranbp3 | Tsc22d2 | Sqle |
| Dlgap4 | Map3k1 | Gpr37 |
| Rps5 | Abcb7 | Ptpn4 |
| Gnb2 | Ranbp2 | Entpd7 |
| Spg7 | Scn3a | Hmgcs1 |
| Golga2 | Klhl7 | Zfp948 |
| Rab11b | Rlim | Homer1 |
| Vps4a | Tra2a | Pcdh9 |
| Kctd13 | Fgd6 | Pcdh20 |
| Morn4 | Ago2 | Tiparp |
| Keap1 | Zfx | Spred1 |
| Rnf220 | Cert1 | Junb |
| Vps52 | Unc80 | Pgap1 |
| Gsk3a | Fem1c | Uba6 |
| Mapk8ip1 | Smad1 | Nedd9 |
| Ssu72 | Osgin2 | Utp14b |
| Actr1a | Pcdh7 | Scn1a |
| Rbm42 | Cntnap5a | Zdbf2 |
| Os9 | Zfp275 | Nkain3 |
| Oscp1 | Col25a1 | Stxbp5l |
| Wbp2 | Snapc1 | Dusp1 |
| Aamp | Kmt2c | Pcdhga10 |
| Psmb2 | Stag2 | Xkr4 |
| Yipf3 | Ncam2 | Pcdhga7 |
| Fzr1 | Hipk1 | Crispld1 |
| Opa3 | Pcm1 | Fzd1 |
| Ssbp4 | Cpeb2 | Map3k9 |
| Ddx54 | Ythdc2 | Klhl11 |
| Arhgdia | Rbm12 | Spry4 |
| Agpat1 | Pde7b | Dusp6 |

| | | |
|---|---|---|
| Map3k10 | Npas3 | Plk3 |
| Arf5 | Gja1 | Sgk1 |
| Arl8a | Sacm1l | Rel |
| Eif2ak1 | Yes1 | Pcsk1 |
| | Nedd9 | Adamts4 |
| | Tmx4 | Il36g |
| | Cacnb4 | Gm32687 |
| | Tbc1d12 | Gpr3 |
| | Vstm2a | Nr4a3 |
| | Slc4a10 | Egr4 |
| | Tmem117 | Sik1 |
| | Fam107b | Trib1 |
| | Bmpr2 | Tent4a |
| | Tmem161b | Atp7a |
| | Rc3h1 | Plekhf1 |
| | Idi1 | Tent5a |
| | Wdfy1 | Egr1 |
| | Pcmtd2 | Nr4a1 |
| | Zfp788 | Fosb |
| | Zfp451 | Arc |
| | Mga | Npas4 |
| | Dnm3 | Fos |
| | Tmx1 | |
| | Shprh | |
| | 1810013L24Rik | |
| | Zfp655 | |
| | Lonrf3 | |
| | Ampd3 | |
| | Tmem74 | |
| | Sntb2 | |
| | Derl2 | |
| | Sptssa | |
| | Atrx | |
| | Srsf10 | |
| | Vps13a | |
| | Id4 | |
| | Esco1 | |
| | Dennd1b | |
| | Tmem38b | |
| | Fndc3a | |
| | Mfsd14a | |
| | Lin7c | |
| | Midn | |
| | Tmed7 | |

Otud4
Purb
Rbm26
Lypla1
Tmem33
Trib2
Edil3
Gria2
Zbtb44
Mfsd14b
Lgalsl
Arfip1
Hook1
Gm45902
Ero1a
Naf1
Tmem170b
Nipbl
Npat
Usp34
Slc2a13
Igip
Nck1
Ptbp2
Usp9x
Slf2
Srsf1
Aqp4
Lig4
Zbtb34
Matr3
Sgk1
Atp2b4
Zfp369
Vcpip1
Dram2
Naa15
Tmem165
Rab39b
Ipcef1
Mex3c
Pde7a
Zfp26
Sp3

Srsf3
Far1
Zfp516
Tm9sf2
Khdrbs2
Pdik1l
Ston2
Rps6ka3
Zfp281
Agfg1
Rhobtb3
Zmym2
Ttc14
mt-Nd5
Tasor2
Sp4
Avl9
Ccdc88a
Rspry1
Kantr
Acer3
Tshz3
Gxylt1
Rab9b
Cfap300
Per2
Msl2
Zfp748
Tet1
Rp2
Yme1l1
Rb1cc1
Lamp2
Ids
Pros1
Tnik
Zbtb11
Mbnl1
Ccnc
Dzip3
Zfp148
Chordc1
Cggbp1
Mib1

Rsbn1l
Cybrd1
Stt3a
Fnip2
Flvcr1
Cfh
Pcdh19
Bcl6
B3galt2
Rock1
Cbfb
C1ql3
Inpp4b
Rfx7
Dio2
Amot
Pcdh17
Xpo4
Mindy2
Ubxn7
Tead1
Utp14b
Srsf7
Pak3
Itgav
Adra1a
Bmpr1b
Tgfbr1
Pdp1
Ints8
Chic1
Arid5b
Pcf11
Fam135b
Thoc2
Mfap3
Ahr
Cdkn1a
Phip
Calb1
Egfr
Creb1
Marcks
Rnft1

Ccdc85a
Med7
Bicd1
Arid4b
Mier3
Vcam1
Mier1
Rfx3
Rora
Nufip2
Rif1
Epm2aip1
Ints6
Chpt1
Cnot6l
Jade3
Cd164
Ranbp6
Pura
Dpy19l4
Larp4
Sec24a
Atp13a3
Dmxl1
Glra2
Zkscan8
Galnt13
Qki
Acvr2a
Xiap
Cdc73
Dact1
Arxes2
Cpeb4
Tbc1d8b
Smc2
Cul4b
Lnpep
Rc3h2
Zfp397
Map3k2
Sacs
Cldn34c1
Zfp770

Pou2f1
Zfp871
Ubn2
Hipk3
Pank3
Osbpl8
Gca
Zfp850
Gm10033
Zbtb41
Spopl
Aasdhppt
Diaph2
Ecm2
Zfp518a
Mospd2
Dok6
Ctdspl2
Grm5
Nxt2
Xpo1
Fnip1
Ap1s2
Agmo
Prpf39
Dr1
Kbtbd7
Ppp1cb
Ptar1
Gng5
Pcgf5
Ireb2
Nexmif
Zfp974
Jag1
Necab1
Zfp874b
Socs4
Hspa13
Taok1
Cacna2d1
Pcdhb18
Tlcd4
Rnpc3

Tasor
Arhgap42
Pcdh15
Cav2
Neurod1
Ppp6c
Vma21
Sox11
Dgkh
Ppp1r3d
Zbtb6
Hecw1
Abca5
Tenm1
Ccdc117
Tnpo1
Fosl2
Cep85l
Haus3
Rbm12b1
Zim1
Dcx
Zfp948
Zc3h12c
Hycc2
Rsbn1
Il1r1
Pgap1
Rbm12b2
Ppp4r3b
G2e3
Arl5b
Bmt2
Styx
Slc7a11
Kdm7a
Rimoc1
Zfp800
Rnf138
Med13
Lcorl
Ro60
Rgs17
Stc1

Ube3a
Zbtb33
Gabrg1
Ptbp3
Kitl
Trib1
Arhgap18
Zfp568
Zfp654
Naa30
Il1rapl1
Dusp6
Gmnc
Scai
Gm48348
Zfp966
Zbtb26
Zfp976
Slc5a3
Rnf217
Cpsf6
Itgb8
Nbeal1
Fbxo30
Uba6
Ptpn4
Sik1
Lpar4
Tmem47
Arhgap5
Tmtc3
Homer1
Trp53inp1
Egr4
Tiparp
Brwd3
Zxdb
St8sia4
Atp11c
Nr4a2
Zfp946
Efna5
Hey2
Plagl1

Zdbf2
Spred1
Pcdh11x
Zfp442
Fut9
BC024063
Egr1
Fosb
Ptchd1
Glra3
Fos
