## Supplemental Table 2 for "Aging disrupts transcriptional programs for memory updating in the dorsal hippocampus and reveals a required role for *Tent5a* in reconsolidation"

| No Update | | Update | |
|---|---|---|---|
| Down | Up | Down | Up |
| Ttr | Gprasp1 | Gm14117 | Ncan |
| Clic6 | D630045J12Rik | Gm56451 | Chrm1 |
| Insc | Ppm1f | Gm5417 | Cdc42se1 |
| Ccdc153 | Foxk2 | Ttr | Trak1 |
| Mfrp | Baz1b | Ak7 | Tex2 |
| Kcnj13 | Nup50 | Dnali1 | Ppm1f |
| Drc7 | Nlgn3 | Clic6 | Foxk2 |
| Dnali1 | Ncan | Mfrp | Speg |
| Cldn2 | Wipf2 | Cldn2 | Zbtb4 |
| Ecrg4 | Rbl2 | Ecrg4 | Ddah1 |
| Aqp1 | Cltc | Aqp1 | Sik3 |
| Hydin | Kctd20 | Col8a2 | Zfp523 |
| Mycbpap | Tbc1d14 | Slc39a4 | Atrn |
| Wdr86 | Camsap1 | Dnah11 | Slc6a1 |
| Stoml3 | Trappc11 | Hydin | Hspa5 |
| Col8a1 | Bbs1 | Dnah6 | Brpf3 |
| Lrrc23 | Snrk | Wdr86 | Nup50 |
| Cfap126 | Uggt1 | C3 | Ubqln4 |
| Cdhr4 | Vps26b | Lrrc23 | Plekhm3 |
| Lbp | Arhgef7 | Ppp1r32 | Srrm2 |
| Cfap65 | Vps39 | Cfap161 | Gaa |
| 2410004PC | Capn15 | H2-Aa | Parp1 |
| Slc39a4 | Hpcal4 | Cfap126 | Lmtk3 |
| Scube3 | Zswim8 | 2410004PC | Ntrk2 |
| Tekt1 | Lrrtm1 | H2-Eb1 | Nf2 |
| Oca2 | Ttc17 | Tekt1 | Atp1b2 |
| Pla2g5 | Fbxl20 | Cd74 | Dact3 |
| Fbln7 | Abcc5 | Nkx3-1 | Ipo9 |
| Otos | Mbd5 | Catip | Arhgef18 |
| Tcte1 | Sez6l | Slc43a3 | Cul4a |
| Ppp1r32 | Cul4a | H2-Ab1 | Ncs1 |
| Klk6 | Zeb1 | Osr1 | Stk24 |
| Togaram2 | Setd5 | Dnai2 | Zmym3 |
| Abhd11os | Tab2 | Slc9a3 | Akap1 |
| Cyp2j12 | Elp1 | Odf3b | Dpp8 |
| Foxj1 | Rere | 1110017D1 | Dstyk |
| Ccdc113 | Spring1 | Ccdc60 | Ranbp10 |
| Trpv4 | Chd4 | Cpxm2 | Phf12 |
| Enkur | Acsl3 | Itgbl1 | Snx30 |
| Sla2 | Morc2a | Sntg2 | Glg1 |
| Odf3b | Klhdc10 | Medag | Mast3 |
| Cd24a | Pnma2 | Cabcoco1 | Marf1 |

| | | | |
|---|---|---|---|
| 1110017D1 | Kdm2a | Slco2a1 | Mrtfa |
| Myl4 | Srrm2 | Rspo2 | Tro |
| Cfap46 | Brd4 | Slamf9 | Rptor |
| Cox6b2 | Plekhm1 | Ubxn10 | Pianp |
| Tppp3 | Sec16a | D830030K2 | Clasp1 |
| Pou6f2 | Sin3a | Bst2 | Hgsnat |
| Klk10 | Zfp532 | 1600029O1 | Scn8a |
| Prkcg | Zfp609 | Ccdc102a | Phactr1 |
| Iyd | Phf2 | Cckbr | Prepl |
| Pdlim1 | Smg7 | Arhgap25 | Cnih2 |
| Nkx3-1 | Kdm5b | Cidea | Brd1 |
| Calb2 | Unc79 | Lpar6 | Ctdsp2 |
| Opalin | Eif4ebp2 | Wnt9a | Kdm3b |
| Prkcd | Pogz | Gpr62 | Sort1 |
| Ninj2 | Bms1 | Tigd3 | Plpp3 |
| Crybg2 | Ccdc6 | Tspan17 | Tecpr2 |
| Clec3b | Baz2a | Egf | Supt6 |
| Gli1 | Irf2bpl | Gm48365 | Wwc1 |
| Rarres2 | Foxo3 | Fmo1 | Fam210b |
| Syt9 | Irs2 | Dusp2 | Zfp142 |
| Proca1 | Map1b | Lcmt2 | Cog6 |
| Serpinb8 | Ago1 | Kcnf1 | Vps39 |
| Serpinb1a | Kdm6b | Efna1 | Ralbp1 |
| Cd59a | Gigyf2 | Gprc5c | Zer1 |
| Gm3558 | Ifnar1 | Anxa2 | Usp8 |
| Capsl | Zzef1 | Slc22a4 | Rbfox2 |
| Srpk3 | Epc1 | Nnat | Cx3cl1 |
| Tmem125 | Dag1 | Scgn | Cacnb1 |
| Prss35 | Fbxo42 | Pygl | Lrp1 |
| Bok | Castor2 | 1700029J0 | Prex1 |
| Hs3st4 | Tmem131 | Spint2 | Rab11fip3 |
| Car14 | Ankrd17 | Gm56450 | Slc12a5 |
| Ubxn10 | Dop1a | Asb16 | Ripor1 |
| Cdsn | Dhx9 | Blnk | Ptpn1 |
| Cfap45 | Rprd2 | Mreg | Adnp2 |
| C1ra | Prrc2b | Tmem196 | Etv3 |
| Cpne9 | Akap9 | Nrde2 | Dgkz |
| Fxyd7 | Ice1 | Spock3 | Fbxl16 |
| Twist2 | Zmiz1 | Ncald | Syngap1 |
| Pdlim2 | C2cd3 | Car10 | Rundc1 |
| Adamts8 | Plekhm3 | Prdx4 | Rai1 |
| Klhdc8a | Cdan1 | Nudt22 | Fbxo42 |
| Cckbr | Arhgef12 | Ak5 | Agap2 |
| Pcolce2 | Iqsec2 | Pet100 | Nr1d1 |

| | | | |
|---|---|---|---|
| Pcsk4 | Sos2 | Lmo4 | Iqsec2 |
| Plpp2 | Lonrf2 | Mctp1 | Arhgap39 |
| Arsg | Phka2 | Lhx9 | Mllt1 |
| Hapln2 | Polr1b | AA414768 | Dusp8 |
| Galnt6 | Epg5 | Klhdc9 | Efna3 |
| Carns1 | Ncor1 | Nmral1 | Brd4 |
| Ccdc102a | Ubr5 | Ppcs | Dag1 |
| Arhgap22 | Ppp1r12b | Acot1 | Tnrc6a |
| Pvalb | Smcr8 | Tagln3 | Anks1b |
| Tspan17 | Fam53b | Ogfrl1 | Magi2 |
| Medag | Tro | Wdr5b | Gigyf2 |
| Gpr62 | Tenm2 | Fabp7 | Kcnab2 |
| Stmn2 | Arhgap21 | Fam183b | Camsap1 |
| Pllp | Ehmt1 | Acy1 | Flt1 |
| Anxa2 | Dido1 | Zswim7 | Hif1an |
| Rspo2 | Brd1 | Neto1 | Ppp1r9b |
| D830030K2 | Cramp1 | S100a6 | Ccdc6 |
| Arhgdig | Lrp1 | Slc26a8 | Dlgap3 |
| Nrip3 | Gldc | Mpp3 | Ubr2 |
| Sfrp4 | Hcfc1 | Sstr3 | Irf2bpl |
| Stac2 | Edrf1 | Polg2 | Snrk |
| Garin5a | Ksr2 | Hyal1 | Tsc1 |
| Sag | Stk24 | Tm4sf1 | Edrf1 |
| Apod | Tpr | Nemp2 | Lrrtm1 |
| Chgb | Scaf4 | Galnt14 | Baz2a |
| B3gat2 | Ttc28 | Anxa4 | Neurl1b |
| Gm5628 | Pde4dip | Tcf7 | Castor2 |
| Adssl1 | Ankrd50 | Oard1 | Crkl |
| Lsm11 | Nsd3 | Cebpzos | Fbrsl1 |
| Bst2 | Med20 | Arhgap15 | Nwd1 |
| Cyp26b1 | Gpatch2l | Sdhaf1 | Gramd1b |
| Arhgap25 | Trmt1l | Snca | Usp21 |
| Prkn | Btbd8 | Lrrc55 | Arel1 |
| Cabcoco1 | Igf2r | Tmeff2 | Ctnnd2 |
| Cd109 | Ubr2 | Smyd2 | Sel1l |
| Fstl3 | Ski | Cd83 | Abitram |
| Sec14l5 | Mon2 | Usp2 | Jph3 |
| Il34 | Ank3 | Pkib | Kdm5b |
| S1pr5 | Kmt2b | Pcdhgc4 | Eif4ebp2 |
| Il33 | Slc35e2 | Clybl | Tbkbp1 |
| 1600029O1 | Chtf8 | Nrn1 | Alkbh5 |
| Ppfibp2 | Nav1 | Gtf2f2 | Sec16a |
| 2310030G0 | Fam43b | Anxa11 | Scaf4 |
| Arrdc2 | Flcn | 3110082I1 | Unc79 |

| | | | |
|---|---|---|---|
| Fmo1 | Anapc1 | Gyg | Phf3 |
| Lpl | Nrcam | Frat1 | Tcf20 |
| Fadd | Atp2b3 | Nudt2 | Man2a2 |
| Ngf | Ckap5 | Necab3 | Dip2c |
| Hyi | Setdb1 | Cdo1 | Specc1l |
| Heyl | Ipmk | Emc9 | Kmt2b |
| 4930523C( | Smarcal1 | Snrnp25 | Grin1 |
| Adora2a | Plxna4 | Trappc1 | Bahd1 |
| Plaat3 | Huwe1 | Pald1 | Jag2 |
| Homer3 | Ep400 | Tipin | Smcr8 |
| Crip1 | Hif1an | Prl8a1 | Ksr1 |
| Zap70 | Zmym3 | Xpa | Pde2a |
| Qdpr | Gan | Cd200 | Fry |
| Slamf1 | Lrp4 | Blmh | Sbf2 |
| Spock1 | Arntl | Amdhd2 | Rcor1 |
| Nnat | Caln1 | Aifm3 | Fam234b |
| Phlda3 | Eaf1 | Sh3gl3 | Snph |
| Krt73 | Taf4 | Atp6v0e2 | Dclk2 |
| Gpc1 | Crnkl1 | Tcta | Camkv |
| Pde8a | Dip2c | Spata33 | Slc25a25 |
| Tmem160 | Stxbp5 | Gpr83 | Agrn |
| Snrnp25 | Adgra2 | Cutc | Cacnb2 |
| Atp5e | Ncoa1 | Slc29a1 | Shank2 |
| Ankrd9 | Dicer1 | Snapc5 | Bms1 |
| Cdpf1 | Chd2 | Inha | S1pr1 |
| Avpi1 | Sec22c | Selenoh | Ice1 |
| Psme2 | Cep162 | Klhl33 | Gan |
| Pet100 | Aqr | Fhip1a | Plxna2 |
| Nhp2 | Med1 | Mcts2 | Snx27 |
| Atp5k | Foxk1 | Tafa2 | Elk4 |
| Prdx4 | Auts2 | Tango2 | Itga6 |
| Pcdhgc4 | Ubn1 | Tgfa | Ylpm1 |
| Morn2 | Zfp598 | Slc48a1 | Smg7 |
| Fhl2 | Myorg | Dcxr | Atxn1l |
| Ostf1 | Plagl2 | Smoc2 | Prrc2c |
| Prob1 | Slitrk5 | Acyp2 | Cramp1 |
| Sult1a1 | Usp8 | Kctd4 | Lrba |
| Cidea | Cdh8 | Pcp4 | Cds2 |
| Nectin4 | Gpam | Rbm11 | Chd3 |
| Cpne4 | Socs7 | Sgf29 | Gabbr1 |
| Dusp26 | Xpo7 | Tmem86a | Nadk |
| Trnp1 | S1pr1 | Uap1l1 | Akap10 |
| S100a6 | Mpdz | Dnaaf4 | Galnt16 |
| Ufsp1 | Sike1 | Morn2 | Cdan1 |

| | | | |
|---|---|---|---|
| Gstt1 | Sbno1 | Zdhhc22 | Tub |
| Stmn1 | Ptprj | Ywhah | Chpf |
| Calhm5 | Crebbp | Tbc1d7 | Gpatch2 |
| Fhit | Tsc1 | Doc2a | Irs2 |
| Itpk1 | Gpatch2 | Grb14 | Soat1 |
| Trpm4 | Bicra | Tmem160 | Ltbp4 |
| Ly6e | Trio | Supt3 | C2cd3 |
| Gng3 | Slk | Glrx | Bod1l |
| Uchl1 | Kansl1 | Macrod1 | Slco1c1 |
| Rassf5 | Baz2b | Dnajb6 | Utp20 |
| Nlrc3 | B4galt5 | Mdp1 | Med14 |
| Gpsm3 | Sec24b | Paip2 | Zfp532 |
| Ldb3 | Per3 | Atp6v1g1 | Myt1l |
| Lynx1 | Herc2 | Tmed9 | Ifnar1 |
| Gamt | Osbpl6 | Srp9 | Ncor2 |
| Gprc5c | Dnmbp | Gsr | Myorg |
| Elavl2 | Dennd4a | Tceal6 | Nsd1 |
| E2f1 | Adgrf5 | Srsf5 | Mon2 |
| Cabp1 | Tnks | Mocs2 | Spata2l |
| ENSMUSG( | Ap1g1 | Nutf2 | Nol4 |
| Lpar6 | Zfp318 | Vdac3 | Gldc |
| Osbpl1a | Zfp142 | Cryzl1 | Trmt1l |
| Acot7 | Wdr3 | Eif1b | Pogz |
| Unc13d | Rnf24 | Tab1 | Pom121 |
| Auh | Txnrd1 | Lgi1 | Mindy2 |
| Ndufa1 | Nwd1 | Znhit2 | Ksr2 |
| Atp5mpl | Notch2 | Manbal | Esyt2 |
| Cnpy2 | Diaph1 | Cisd1 | Ankrd11 |
| Ndufaf8 | Nedd4 | Srd5a3 | Tmem131 |
| Prxl2b | Snx27 | Serpinb6a | Ppp1r15b |
| Znhit2 | Strn3 | Pfdn2 | Epg5 |
| Cox7c | Prex1 | Ndufa1 | Pde4dip |
| Nrde2 | Msh6 | Cyb5a | Arhgef12 |
| Haghl | Rcan3 | Atosa | Kdm5c |
| Ndufs6 | Lrrn3 | Hdgfl2 | Srcap |
| Mrps33 | Dusp8 | Rassf3 | Stox2 |
| Cops9 | Lmnb2 | Nt5c | Dpy19l3 |
| Pdcd5 | Tcf20 | St3gal5 | Kctd1 |
| Gyg | Prr12 | B4gat1 | Map2 |
| Ndufb8 | Synpo | Ppib | Spring1 |
| Fis1 | Lig3 | Auh | Tcof1 |
| Cox7b | Lrch3 | Nmnat2 | Zfp280b |
| Smim11 | Fbrsl1 | Akr1a1 | Kmt2e |
| Rnh1 | Lbr | Fam241b | Tjp1 |

| | | | |
|---|---|---|---|
| Ndufs5 | Akt3 | Gpr162 | Dgkg |
| Snapc5 | Eml4 | Cndp2 | Fasn |
| Phyh | Tanc2 | Atraid | Wasf3 |
| 1110065P2 | Ncbp3 | Ndufb9 | Rnf145 |
| Ndufb9 | Marf1 | Psmb4 | Pkd1 |
| Mrpl58 | Tbcel | Pccb | Mical2 |
| Rpl36 | Vwa8 | Pfdn1 | Hcfc1 |
| Myl6 | Utp20 | Prxl2b | Mex3d |
| Rabac1 | Nup210 | Ccdc107 | Med13l |
| Nudt2 | Trip11 | Atp5mpl | Lpin1 |
| Cox4i1 | Zfp334 | Zmat2 | Crebbp |
| Tmem256 | Dnajc13 | Cox7b | Ptpn9 |
| Znrd2 | Smc3 | Rtca | Golgb1 |
| Ccs | Ncoa6 | Gstm7 | Atf7ip |
| Ndufa2 | Alkbh5 | Hint1 | Cdk5r2 |
| Vsnl1 | Zfp280b | Cldn5 | Tti1 |
| Stmn3 | Ssh2 | Retreg1 | Tesk1 |
| Ndufa13 | Chd7 | Ndufb3 | Notch1 |
| Msrb2 | Slitrk3 | Nicn1 | Adgrb2 |
| Smdt1 | Ankfy1 | Gcsh | Nup98 |
| Nudc | Eps8 | Acadm | Psd |
| Ubl5 | Wdcp | Phyh | Kdm4b |
| Rbm11 | Etv3 | Parl | Rasgrf1 |
| Tbcb | Gspt2 | Tmem135 | Igdcc4 |
| Cebpzos | Myo9a | Faim | Socs7 |
| Pfdn1 | Adcy6 | Mrpl27 | Tmf1 |
| Npepl1 | Nalcn | Rit2 | Chd5 |
| Tmem86a | Son | Commd1 | Plekhm1 |
| Atp5j2 | Pik3r3 | Ndufb8 | Celf5 |
| Rpl38 | Rad54l2 | Nt5c3b | Zfyve26 |
| Uqcrq | Pcid2 | Rnf112 | Map1a |
| Drap1 | Cnot4 | Matk | Brd3 |
| Gfod2 | Chd3 | Snx7 | Spred3 |
| Gmpr2 | Wars2 | 2210016L2 | Kdm2b |
| Ndufa3 | Fhip1b | Nsmce2 | Cnot1 |
| D8Ertd738 | Usp47 | Gstz1 | Fbxl14 |
| Arl6ip4 | Brd3 | Pex16 | Gpam |
| Fkbp2 | Serac1 | Dhdds | Otub2 |
| Uqcrh | Pde10a | Stard7 | Camk2a |
| Hcfc1r1 | Arhgap32 | Rhbdl1 | Ubn1 |
| Pin4 | Jmy | Ndufs6 | Med1 |
| Atp6v1f | Proser1 | Nudt16 | Taf2 |
| Tmed3 | Atf7 | Shisa4 | Stk4 |
| Paxx | Bbx | R3hcc1 | Wdcp |

| | | | |
|---|---|---|---|
| Tmem9 | Mcl1 | Prune2 | Uck2 |
| Ndufb7 | Sfpq | Slc25a28 | Ank3 |
| Nudt22 | Dpp8 | Thoc7 | Cacng8 |
| Cuta | Spata2l | Tsr3 | Las1l |
| Edf1 | Kdm5c | Otulin | Wwc2 |
| Sdhaf1 | Dstyk | Surf1 | Kat6a |
| Cox6b1 | Vldlr | Selenof | Hmgcr |
| Pttg1 | Cacna1h | Use1 | Zhx3 |
| Pebp1 | Add2 | Anp32b | Agfg2 |
| Selenow | Pcdhgc3 | Slc35b1 | Tnr |
| Slc48a1 | Noc3l | Dusp14 | Nav1 |
| Pnpo | Pcdhga12 | Rpl22 | Drp2 |
| Atp5o | Ankrd11 | Imp3 | Usp42 |
| Mgst3 | Mphosph8 | Psmd4 | Zmiz1 |
| Mocs1 | Slit1 | Zcrb1 | Bbx |
| Ap2s1 | Atrn | Dlat | Mecp2 |
| Lcmt2 | Rbm33 | Mcu | Macf1 |
| Tango2 | Stk4 | Gstm4 | Btaf1 |
| Rhou | Rapgefl1 | Hspa2 | Pcdhgc3 |
| Vamp1 | Herc1 | Psme1 | Mtf1 |
| Oard1 | Dnmt3a | Mrps33 | Kansl1 |
| H2-T22 | Pom121 | Kcng2 | Kdm5a |
| Anapc13 | Pcgf3 | Psmc5 | Dlg2 |
| Bpgm | Cplane1 | Mrpl17 | Ago3 |
| Mad2l2 | Slc39a14 | Aldh9a1 | Ski |
| Mcee | Pcdhgc5 | Cfap74 | Rprd2 |
| Sem1 | Tlnrd1 | Sod1 | Pde10a |
| Pacsin3 | Dtx4 | Comt | Zfp592 |
| Gstp1 | Pcnx | Ctsl | Eml6 |
| Plekho1 | Gpd2 | Sem1 | Klf10 |
| Clybl | Slc12a5 | Syt13 | Cnksr2 |
| Atpif1 | Slc12a6 | Hsbp1 | Zfta |
| Tomm6 | Plxna2 | Snx24 | Extl1 |
| Bbln | Mapk7 | Plpp1 | Dlg4 |
| Klhdc9 | Zfp12 | Echdc1 | Crnkl1 |
| Sh3gl3 | Rsf1 | Hes6 | Naf1 |
| Nedd8 | Nup205 | Tmem234 | Hmgxb3 |
| Ndufa11 | Hectd4 | Tmem25 | Panx2 |
| Oxld1 | Tshz1 | Cmas | Trrap |
| Rab4b | Dpy19l3 | Tmem38a | Synpo |
| Ndufa7 | Nf1 | Rpl21 | Ncbp3 |
| Gpx4 | Tub | Mtarc2 | Khdc4 |
| St8sia6 | Rai1 | Txn1 | Ap2b1 |
| Fbxo2 | Dnaaf10 | Pgrmc2 | Ints12 |

| | | | |
|---|---|---|---|
| Vps28 | Pdpr | Mrps34 | Nav2 |
| Cisd3 | Ccar1 | Mrpl36 | Adgra2 |
| Psme1 | Usp15 | Atp5e | Trappc8 |
| Idnk | Pag1 | N6amt1 | Npat |
| Rnaseh2c | Bhlhe41 | Rpap3 | Mturn |
| Septin8 | Ammecr1l | Nedd8 | Ptch1 |
| Fth1 | Bcl9l | Ap1m1 | Ncor1 |
| Uqcr10 | Jph4 | Ppia | Kctd21 |
| Uqcr11 | Sptb | Kcns1 | Apc2 |
| Yipf2 | Dlg2 | Nck2 | Celsr3 |
| Trnau1ap | Map3k13 | Rab24 | Adgrf5 |
| Pepd | Rlf | Kxd1 | Abcb1a |
| Gng13 | Zfp180 | Fndc10 | Dennd4a |
| Sult4a1 | Magi2 | Dynlt1b | Pikfyve |
| Cryzl2 | Ddx18 | Qtrt1 | Slc35e2 |
| Aig1 | Bicral | Rpl38 | Vgll4 |
| Msrb1 | Zzz3 | 1110032A0 | Grin2c |
| Ggact | Brpf3 | Nfkbil1 | Ccdc85c |
| Map1lc3a | Rbm4b | Cirbp | Shisa6 |
| Rpl41 | Cacna1e | Ptprk | Ago1 |
| Romo1 | Nalf2 | Txnl4b | Kcnk12 |
| Ndufv3 | Med14 | Micos13 | Bicral |
| Tubb4b | Rpap1 | Mycl | Pcdh20 |
| Dynlt1b | Upf2 | Nek6 | Cnot4 |
| Mt3 | Sipa1l3 | Pced1b | Klf15 |
| Ccdc28a | Glg1 | Fabp5 | Tspoap1 |
| Tuba1b | Heatr1 | Psme2 | Ssh2 |
| Tcta | Fbxo41 | Tbca | Proser1 |
| Lage3 | Zfp608 | Cfdp1 | Nf1 |
| Gpank1 | Zfp592 | Cetn3 | Atp2b3 |
| Mien1 | Tti1 | Sdhaf4 | Ccnt1 |
| C1ql1 | Rnf38 | 1110065P2 | Smurf2 |
| Spata33 | Mtcl1 | Pfdn6 | Zfp365 |
| Ilvbl | Cers6 | Upf3a | Nsd3 |
| Mrpl54 | Trpm3 | Cmpk2 | Ncoa2 |
| Cdkn1a | Selenoi | Naglu | Ep400 |
| Tuba1a | Gna13 | Atp6v1e1 | Tbl1x |
| Syde1 | Paqr9 | Wrap73 | Lmnb2 |
| Nudt18 | Ttll4 | Avpi1 | Trip12 |
| Psmb10 | Mrtfb | Tpt1 | Plec |
| Dap | Ubr4 | Psmb1 | Togaram1 |
| Fn3k | Zbtb21 | Gstm5 | Soga3 |
| Wwox | Cds2 | Prxl2a | Cdk13 |
| Grb14 | Dmxl2 | Ndufs3 | Prrc2b |

| | | | |
|---|---|---|---|
| Srp9 | Cacna1b | Rhoa | Slc1a2 |
| Atp6v1g1 | Tanc1 | Glo1 | Unc13b |
| Pfdn2 | Fem1b | Rabggtb | Pptc7 |
| 5730409EC | Mphosph9 | Psmd13 | Zfp319 |
| Cops6 | Cc2d2a | Pithd1 | Foxg1 |
| Tpt1 | Atp8a1 | Nipsnap1 | Anapc1 |
| Acsl5 | Crlf3 | 4933434E2 | Nin |
| Dmac1 | Utp4 | Ank | Taf4 |
| Tceal6 | Stk35 | Nipsnap2 | Zfp866 |
| Ndufa6 | L2hgdh | Bdh1 | Diaph1 |
| Gm2000 | Dot1l | Psmc2 | Pkn2 |
| Anp32b | Fam91a1 | Mcfd2 | Prrc2a |
| Tmem234 | Pik3ca | Vdac1 | Dtx4 |
| Zcrb1 | Atg14 | Tmem59 | Kdm1b |
| Rtca | Stard13 | Ywhab | Stk40 |
| Gstm5 | Kdm3b | Ilf2 | Trim37 |
| Ndufb3 | Zfp445 | Slc25a4 | Snd1 |
| Parl | Gspt1 | Sdhaf2 | Helz |
| Tstd3 | Nos1 | Tmx2 | Edem3 |
| Mrps36 | Pitpnc1 | Dazap2 | Hipk1 |
| Manbal | Polk | Rraga | Caskin1 |
| Nkiras1 | Znrf3 | Dact2 | Ddx19b |
| Surf1 | Tspoap1 | Atp6v1d | Hycc1 |
| Atp6v0b | Tstd2 | Clta | Gpr176 |
| Pex16 | Prdm2 | Acsl5 | Zbtb16 |
| Rps5 | Zswim5 | Sdhd | Nalf2 |
| Eif1b | Lamc1 | Acbd6 | Enah |
| Atp5h | Gse1 | Cdc42 | Tom1l1 |
| Ndufs3 | P4ha1 | Ankmy2 | Camsap2 |
| Ranbp1 | Spred2 | Becn1 | Gigyf1 |
| Mocs2 | Nup98 | Cct2 | Gm48368 |
| Ndufc1 | Abitram | Dhrs7 | Jmjd1c |
| Mpnd | Prickle1 | Arl6ip5 | Rasal1 |
| Psmb4 | Rhobtb2 | 5730409EC | Ralgps1 |
| Tmed9 | Lrba | Vdac2 | Smad4 |
| Eif2b1 | Dnajb5 | Fstl1 | Maml1 |
| Atp5l | Zfyve26 | Cyth2 | Mras |
| Oaz1 | Igdcc4 | Oaz1 | Myh9 |
| Fbl | Rcor1 | Klhl26 | Ing5 |
| Use1 | Slx4 | Myl12b | Lix1l |
| Mrto4 | Arid1a | Ndufs2 | Spred2 |
| Ndufb11 | Csmd2 | Suclg1 | Kdm2a |
| Gpr162 | Med12 | Prdx1 | Kdm6b |
| Nme2 | Ylpm1 | Rab7 | Sptbn2 |

| | | | |
|---|---|---|---|
| Snx15 | Mdn1 | Gphn | Trio |
| Akr1a1 | Hectd1 | Keap1 | Gpt2 |
| Rpl23 | Klf12 | Saraf | Map3k13 |
| Rps29 | Bod1l | Ywhaz | Fam43b |
| Nubp2 | Trrap | Atp5l | Gfod1 |
| Bckdha | Tnrc6a | Aph1a | Sipa1l3 |
| Rpl37 | Prrc2c | Atp5g3 | Prdm2 |
| Gde1 | Kdm3a | Fam216a | Zswim5 |
| BC029722 | Phf3 | Zfp706 | Slx4 |
| Msra | Ryr2 | Gnptg | Bicra |
| Emc4 | Bcorl1 | Rps20 | Ppp1r3c |
| Ddx25 | Kcnk12 | Cct6a | Srf |
| Timm8b | Sbf2 | Cox5a | Samd4b |
| B4gat1 | Bptf | | Arid1a |
| Uqcc2 | Zfhx2 | | Osbpl6 |
| Dact2 | Nsd1 | | Dido1 |
| Timm13 | Hivep3 | | Heatr5b |
| Aspscr1 | Akap10 | | Fgd6 |
| Rpl26 | Nfya | | Tenm2 |
| Rpl24 | Pkd1 | | Exoc6b |
| Park7 | Setd1b | | Ptprb |
| Psmb6 | Tnrc6c | | Zbtb11 |
| Suclg1 | Dcaf1 | | Sorl1 |
| Cdk5 | Tnr | | Ncoa1 |
| Tmx2 | Birc6 | | Dlgap2 |
| Serpinb6a | Fam120c | | Prickle1 |
| Nme1 | Plekha8 | | Nfya |
| Rpl11 | Per1 | | Lamc1 |
| Ddrgk1 | Ddi2 | | Gse1 |
| Pgam1 | Mycbp2 | | Creb3l2 |
| Rps14 | Atg2b | | Pprc1 |
| Ndufs2 | Taf2 | | Lrp4 |
| Rab24 | Smad1 | | Ahdc1 |
| Rpl36a | Sorbs1 | | Slit3 |
| Mrpl27 | Dcp1a | | Cmtm4 |
| Gpr180 | Tnrc18 | | Arnt2 |
| Mrpl42 | Ahdc1 | | Zfp598 |
| Rps28 | Aff4 | | Bcorl1 |
| Cd200 | Khdc4 | | Zfp710 |
| Mrpl28 | Virma | | Lmtk2 |
| Cuedc2 | Ptch1 | | Mettl16 |
| Sdhc | Kdm5a | | Zcchc14 |
| Larp6 | Tjp1 | | Prr14l |
| Vdac1 | Atf7ip | | Arhgap33 |

| | | |
|---|---|---|
| Ndufa4 | Jmjd1c | Dcaf1 |
| Tmem242 | Slit3 | Bcor |
| Rps25 | Zcchc14 | Rhobtb2 |
| Ndufb2 | Cnot1 | Tnrc6c |
| Hint1 | Frmd4a | Bcl9l |
| Cox5a | Peg3 | Setd1b |
| Psmc5 | Spata13 | Tmem38b |
| Naa12 | Zfp236 | N4bp1 |
| Acaa1a | Cmtm4 | Tnrc18 |
| Eefsec | Vgll4 | Phf8 |
| Psmb1 | Heatr5b | Frmd4a |
| Vps51 | Tet3 | Phf13 |
| Emc9 | Med13l | Dnajc13 |
| Rpl32 | Sbk1 | Bptf |
| Cox8a | Csmd3 | Kmt2a |
| Rnf220 | Mecp2 | Chd1 |
| Tle5 | Spry4 | Arhgap32 |
| Naxe | Pikfyve | Septin9 |
| Eif3k | Dock7 | Med23 |
| Matk | Prr14l | Adcy9 |
| Ndufa5 | N4bp1 | Sec14l1 |
| Coa3 | Kmt2e | Zfp180 |
| R3hcc1 | Ago3 | Asap1 |
| Commd1 | Pcm1 | Cdk12 |
| Txndc17 | Ahctf1 | Pou6f1 |
| Psmb5 | Utp14b | Kcnb1 |
| Cox5b | Celsr3 | Csmd2 |
| Slc39a7 | Atm | Rnf24 |
| Rpl21 | Setx | Ptpn12 |
| Swi5 | Pprc1 | Zfp236 |
| Nutf2 | Ankhd1 | Med12 |
| Mdp1 | Stox2 | Caln1 |
| Rps27 | Zcchc2 | Peg3 |
| Fam241b | Lrrc7 | Jph4 |
| Mdh1 | Kat6a | Sptb |
| Cope | Zbtb20 | Pakap |
| Srsf5 | Mn1 | Pnkd |
| Sharpin | Junb | B3gat1 |
| Tm2d3 | Lrp1b | Cbl |
| Cox7a2 | Elfn2 | Tek |
| Sdhb | Nbea | Ttc39b |
| Cisd1 | Bltp1 | Ddx18 |
| Fabp5 | Setd2 | Tnrc6b |
| Ppia | Zfp516 | Kcnn2 |

| | | |
|---|---|---|
| Ddt | Prdm11 | Aff4 |
| Pdss2 | Btaf1 | Vstm2l |
| Pmm1 | Zfp831 | Ccdc117 |
| Cdo1 | Nin | Kmt2d |
| Sec11c | Kcnj6 | Zmat3 |
| Upf3a | Stam2 | Zfp467 |
| Dctn5 | Otub2 | Rbm12b1 |
| Pam16 | Dync2h1 | Kcnj6 |
| Iscu | Slc1a2 | Gm1043 |
| Rpl12 | Tek | Arhgap26 |
| Pfdn6 | Arid2 | Zfyve16 |
| Psmb7 | Ash1l | Rims4 |
| Selenok | Zfp638 | Tfdp2 |
| Cycs | Zfp866 | Shank3 |
| Lin7b | Mtr | Crim1 |
| Ube2s | Sesn3 | Mia3 |
| Tmem59l | Usp27x | Fam120c |
| Fndc10 | Insr | Dusp4 |
| Tmem258 | Usp37 | Cep350 |
| Elof1 | Zbtb11 | Zbtb21 |
| Rnf25 | Dusp4 | Aff1 |
| Capns1 | Sec24d | Smad1 |
| Natd1 | Ptprb | Nufip2 |
| Rps19 | Pdp1 | Mideas |
| Cyb5a | Cep295 | Rab3c |
| Idh3g | Nav2 | Usp31 |
| Ubxn1 | Togaram1 | Ash1l |
| 2210016L2 | 9930021J03Rik | Ago2 |
| Nicn1 | Usp24 | Pde5a |
| Tusc2 | Notch1 | Zfp169 |
| Fam3a | Dock3 | Plagl2 |
| Micos10 | Egr3 | Zfp804a |
| Mif | Pum1 | D430019H16Rik |
| Aarsd1 | Sall2 | Tsc22d3 |
| Cox6a1 | Sphkap | Ank1 |
| Ube2m | Chd1 | Zc3h12c |
| Stub1 | Ptpn12 | Tet3 |
| Grcc10 | Pdpk1 | Marchf4 |
| Arf5 | Prkdc | Egr3 |
| Atp5d | Soga1 | Ptcd1 |
| Rnf5 | D430019H16Rik | Lmo1 |
| Maea | Zfp667 | Zfp704 |
| Scn1b | Tmem38b | Ddn |
| Snx7 | Scn2a | Shisa7 |

| | | |
|---|---|---|
| Rplp2 | Lifr | Intu |
| Mpst | Med12l | Usp27x |
| Fkbp8 | Ccdc117 | Nup153 |
| Rpl34 | Itga6 | Ern1 |
| Adprh | Pigg | Dcp1a |
| Nenf | Sipa1l1 | Ampd3 |
| Serf2 | Sorl1 | Mtcl1 |
| Uba52 | Bdp1 | Zfp516 |
| Tomm7 | Reln | Zfp526 |
| Psmc3 | Dcp2 | Ankrd33b |
| Dad1 | Mex3c | Oprd1 |
| Mrps34 | Trip12 | Fgfr3 |
| Pgk1 | Aff1 | Paqr3 |
| Gtpbp6 | Urb2 | Fosl2 |
| Tmem25 | Kcnb1 | Arid5a |
| Rab3a | Lct | Fat3 |
| Snf8 | Tet1 | Csmd1 |
| Smyd2 | Disp1 | Nos1ap |
| Bsg | Flt1 | Klf2 |
| Dnajc8 | Ncoa2 | Adcy2 |
| Ccdc12 | Ppp1r9a | Cntnap2 |
| Cdkn2d | Pcdhb3 | Slc1a6 |
| Mrpl12 | Med23 | Tmsb10 |
| Mtarc2 | Tut4 | Ccdc120 |
| Zkscan17 | Rnf111 | Otud1 |
| Coq7 | Frmd6 | Ranbp2 |
| Syp | Ercc6 | Ppp1r3g |
| Rad23a | Hrk | Setbp1 |
| Pgrmc2 | Map2 | Mga |
| Selenos | Tnks2 | Trmt44 |
| Surf2 | Lrp6 | Kif21b |
| Rpl31 | Pcdhgb6 | Fgf11 |
| Rps27a | Wnk3 | Arntl |
| Psmc4 | Zfyve16 | Dio2 |
| Hdac11 | Mettl16 | Pcdhga2 |
| Dhrs7 | Ripor2 | Adgrb3 |
| Atp6v0c | Ccdc120 | Sntb2 |
| Cdk10 | Gabrb1 | 9930021J03Rik |
| Rps15 | Phlpp2 | Trib1 |
| Rpl18a | Pvr | Map3k1 |
| Dctn3 | Hace1 | Zkscan16 |
| Fam98c | Usp13 | Frmpd3 |
| Gcsh | Edem3 | Kmt2c |
| Chmp2a | Hipk1 | Mex3c |

| | | |
|---|---|---|
| Rnf187 | Trappc8 | She |
| Mrpl17 | Ntrk3 | Vgf |
| Sdr39u1 | Osbpl11 | Sema6c |
| Ttc27 | Ankrd33b | Pcdhgb7 |
| Tab1 | Atxn7 | Plxnd1 |
| Lias | Tlr3 | Nhsl2 |
| Eif6 | Gm48368 | Ext1 |
| Ccm2 | Unc80 | Mlip |
| Rps17 | Zbtb2 | Zfp551 |
| Tpst2 | Celf1 | Gm42669 |
| Scamp4 | Fhip2a | Elfn2 |
| Rpn1 | Zbtb34 | Jade2 |
| Thy1 | Usf3 | Per1 |
| Anapc11 | Ppm1d | P4ha1 |
| Ndufv2 | Pkn2 | Arid4a |
| Cyc1 | Nanos1 | Osbpl11 |
| Selenoo | Asxl2 | Dab2ip |
| Mrpl20 | Rc3h1 | Dnajb5 |
| Ndufaf5 | Cttnbp2 | Sipa1l1 |
| Cstpp1 | Ttc26 | Vegfa |
| Micos13 | Papolg | Kdm3a |
| Coa8 | Lrrn1 | Lrrc7 |
| Ldhb | Zfp790 | Hivep3 |
| Ssr4 | Dennd5b | Ep300 |
| Cfdp1 | Tet2 | Git1 |
| Msto1 | Mindy2 | Smurf1 |
| Scg5 | Mia3 | Zfp703 |
| Rps16 | Vps13a | Sik2 |
| Supt4a | Wdfy3 | Etv5 |
| Abhd16a | Zfp644 | Stum |
| Golga7b | Slc4a4 | Trpc5 |
| Rps10 | Cdk7 | Urb2 |
| Fau | Fnip2 | Adgrl3 |
| G6pc3 | Pop1 | Slc24a3 |
| Rpl19 | Pcf11 | Foxo6 |
| Glrx | Ankrd12 | Tlnrd1 |
| Sod1 | Smurf2 | Pigg |
| Rrp9 | Pcdh8 | Mn1 |
| Cmpk2 | Peg10 | Hivep2 |
| Sfxn3 | Npat | Tox2 |
| Ankrd24 | Zfp280d | Entrep2 |
| Dctn2 | Top1 | Jcad |
| Chpf2 | Smchd1 | Map4 |
| Ndufv1 | Usp9x | Slf2 |

| | | |
|---|---|---|
| Snca | Cbx5 | Frmd6 |
| Pithd1 | Ryr1 | Cttnbp2nl |
| Mrps26 | Shank1 | Prkcb |
| Mtres1 | Kdm6a | Grin2b |
| Sap18b | Ltn1 | Sh3bp4 |
| Rps24 | Rasal2 | Ryr3 |
| Rps26 | Rb1cc1 | Fosb |
| Arpc3 | Nck1 | Adra2c |
| Ftl1 | Pds5b | Cdh10 |
| Ech1 | Pdzd8 | Bend4 |
| Tpgs1 | Eea1 | Mafk |
| Gabarap | Zc3h12b | Plk2 |
| Ostc | Zfp971 | Kctd15 |
| Bin1 | Lmln | Man2a1 |
| Rabl2 | Spty2d1 | Sh3rf1 |
| Ndufaf3 | Arfgef1 | Tle3 |
| Tmem276 | Spon1 | Gadd45b |
| Cox6c | Zfp382 | Mfap3 |
| Uqcrc1 | Usp31 | Tmem74 |
| Nelfe | Ubr1 | Ints6 |
| Tmem14a | Ahi1 | Tcp11l1 |
| Znhit3 | Ddn | Slc9a7 |
| Basp1 | Cep76 | Trpc4 |
| Comt | Cfap97 | Spon1 |
| Psmd4 | 1810013L24Rik | Spats2l |
| Ttc9b | Egfr | Dnah9 |
| Mcrip1 | Pard3b | Cep85l |
| Slc2a6 | Fat3 | Kctd16 |
| Uqcc3 | Prps1l3 | Sall3 |
| Dhps | Arid4a | Scn4b |
| Nudt19 | Kmt2a | Zfp366 |
| Aamdc | Cdk12 | Spred1 |
| Scand1 | Gfod1 | Serinc2 |
| Thtpa | Pcdhgb7 | Usp25 |
| Cfl1 | Ep300 | Uprt |
| Stmn4 | Spen | Sox21 |
| Asphd1 | Helz | Dusp1 |
| Atp6v1e1 | Fgd6 | Tyro3 |
| Trappc1 | Cbl | Dyrk3 |
| Psmb3 | Cep350 | Fndc1 |
| Slc25a4 | Bhlhe40 | Zfp882 |
| Cox7a2l | Mideas | Plekha7 |
| Dctn6 | Kmt2d | Shb |
| Fbxw7 | Ppp1r3c | Car7 |

| | | |
|---|---|---|
| Psmd13 | Ago2 | Pdzrn4 |
| Rab7 | Tnrc6b | Pcdhgb4 |
| Actb | Csmd1 | Zfp800 |
| Dlat | Vps13c | Wdr44 |
| Sdhaf2 | Vegfa | Pcdhga10 |
| Zcchc18 | Kmt2c | Gm28036 |
| Uqcrfs1 | Bcl6 | Rsl1 |
| Rps3 | Gm1043 | Gimap6 |
| Txn2 | Pcdhga2 | Fzd1 |
| Bccip | Zfp169 | Nhs |
| Gabarapl1 | Ston2 | Grin2a |
| Vdac3 | Dennd4c | Npas2 |
| Arfip2 | Nuak1 | Samd5 |
| Dynll1 | Pcdh20 | Zfp874b |
| Saraf | Rev3l | Zfhx2 |
| Clta | Setbp1 | Nuak1 |
| Naa10 | Zfp704 | Shank1 |
| Becn1 | Ccnt1 | Midn |
| Ctsf | Nup153 | Spen |
| Eef1b2 | Map2k6 | Klhl11 |
| Pdcd6 | Zkscan16 | Galnt9 |
| Mtch1 | Map3k1 | Spry4 |
| Coq2 | Prkd2 | Homer1 |
| Nipsnap1 | Ranbp2 | Cacna1g |
| Eif5a | Oprd1 | Bcl6 |
| Pcmt1 | Bves | Slc24a4 |
| Oaz2 | Adcy1 | Ephb3 |
| Dalrd3 | Adgrl3 | Arid5b |
| Arl3 | Cacna1i | Sik1 |
| Rpl14 | Nhsl2 | Bhlhe40 |
| Keap1 | Arid5b | Rasd1 |
| BC031181 | Gm42669 | Dgki |
| Ngrn | Fosb | Astn2 |
| Alg2 | Nipbl | Epha6 |
| Nars | Sik1 | Pou3f3 |
| Rpl13 | Smc2 | Gabrb2 |
| Ssr2 | Smg1 | Lcor |
| Kdelr1 | Casp8ap2 | Ttc9 |
| Eif2b5 | Fndc3a | Hunk |
| Gnb2 | Btbd7 | Iqgap2 |
| Acat1 | Trmt44 | Camk1g |
| Timm23 | Lonrf3 | Nrip1 |
| | Sik2 | Shc3 |
| | Tnik | Atp2b1 |

Rictor
Grin2b
Prkg1
Zbtb37
Mmp16
Pakap
Foxo6
Bcr
Zc3h12c
Itpr1
Arl5b
Ptprz1
Scn3a
Zfp951
Tmem67
Hmcn1
Dio2
Syt10
Iqgap2
Cul4b
Hcfc2
Pou2f1
Glcci1
Zfp292
Shprh
Crebrf
Bmpr1b
Lyst
Zfp871
Atp13a3
Homer1
Usp34
Pdzd2
Fam135b
Rfx7
Atrx
Sacs
Arid4b
Nexmif
Col25a1
Cntnap5b
Glt8d2
Lgr4
Lypd1
Matn2
Pcdhga1
Foxp1
Dll1
Fam78a
Glt8d2
Rel
Hcn1
Cd34
Dact1
Tmem158
Uba6
Inhbb
Adcy8
Itpka
Gabrg3
Lypd1
Thrb
Cds1
Parm1
Baiap3
Penk
Gab3
Sbk1
Zfpm1
Cbfa2t3
Rara
Bcr
Itpr1
Il36g
Cacna1i
Plekhg1
Zdbf2
Chrm4
Adam12
Asap2
Atp7a
Flrt1
Entpd4
Tbc1d1
Doc2b
Rasl11b
Gucy1a2
Slc8a1

Tsc22d2
Prex2
Zfp369
Trpc5
Fosl2
Npy5r
Adgrl4
Tasor2
Qser1
Zfp266
Aff2
Otud4
Cnot6
Grin2a
Sp4
Skil
Zfp397
Prox1
Wdr44
Kcnk9
Tmppe
Dock10
Chl1
Dock11
Pcdhb11
Gria2
Fnip1
Fndc1
Erbb4
Dll1
Sema5a
Slf2
Per2
Klhl11
Slc24a4
Egr4
Dgki
Slc9a7
Mga
Zfp366
Ints6
Gabrb2
Sox21
Dact1
Cadps2
Zfhx4
Epop
Atg9b
Apold1
Ar
Cpne8
Tmem200a
Rab8b
Espn
Lancl3
Spink8
Tnfrsf25
Cfap100
Per2
Junb
Dusp6
Ppargc1b
Acer2
Sez6
Zfp462
Ccnd1
Pcdhb2
Pex5l
Gpr63
Pakap
Homer2
Cdkl4
Tenm3
Galntl6
Meis2
Tent4a
Crhbp
Efnb2
Egr4
Tnfaip3
Sorcs3
Ccdc88c
Igsf9b
Fbn2
Akr1c18
Egfl6
BC030500
Ror1

Smarcad1
Mfap3
Fat4
Nufip2
Xkr4
Pcdh19
Map3k2
Doc2b
Med13
Pcdhb16
Kif26b
Gucy1a2
Pura
Tmem74
Zdbf2
Nrip1
Zfp874b
Cecr2
1700086L19Rik
Zfp800
Itga4
Thsd7a
Pcdhb21
Dmxl1
Spred1
Htr4
Pcdh17
Ptchd1
Atp7a
Acrbp
Tent4a
Greb1l
Diaph2
Homer2
Rab8b
Sstr1
Acer2
Zim1
Ppargc1b
Zfp882
Kctd16
Cep85l
Lcor
Sntb2

Hs6st3
Lefty1
ENSMUSG00000121307
Wnt2
Tac2
Nr4a2
Sstr4
Depp1
Ctxnd1
Ptpru
Dach1
Pou3f1
Fos
Ano5
Gm19410
Pomc
Tmem252
Chrm5
Egr1
Htr5b
Adgrl2
Wfs1
Gpr161
Arc
Alox12b
Nr4a1
Grem1
Egr2
Gh

Slc16a10
Kcna3
Uprt
Arc
Kdm7a
Ptchd4
Nkain3
Col19a1
Nt5c1a
Hs6st3
Onecut1
Tenm1
Tmtc3
Zfhx4
Cth
Vmn1r207
Cbfa2t3
Pcdhb2
Uba6
Egr1
Nr4a1
Depp1
Or6c208
Tnfrsf25
Sox11
Sytl5
Or1n2
Nr4a2
Fos
Zfp458
Kcng3
H3c14
Ghsr
Gm48035
Vmn1r47
Gm3325
Pcdha8
Egr2
