## Supplemental Table 4 for "Aging disrupts transcriptional programs for memory updating in the dorsal hippocampus and reveals a required role for *Tent5a* in reconsolidation"

Gh
Egr2
Grem1
Alox12b
Gpr161
Adgrl2
Htr5b
Wfs1
Chrm5
Tmem252
Gm19410
Pomc
Ano5
Depp1
Pou3f1
Sstr4
Ctxnd1
Dach1
Nr4a2
Ptpru
Tac2
Lefty1
Wnt2
Hs6st3
ENSMUSG00000121307
Egfl6
BC030500
Ccdc88c
Fbn2
Tnfaip3
Sorcs3
Igsf9b
Ror1
Akr1c18
Meis2
Zfp462
Galntl6
Pakap
Per2
Cdkl4
Pex5l
Sez6
Pcdhb2
Homer2

Acer2
Gpr63
Ccnd1
Tenm3
Ppargc1b
Crhbp
Efnb2
Zfpm1
Flrt1
Spink8
Zfhx4
Tmem200a
Doc2b
Cbfa2t3
Cacna1i
Slc8a1
Asap2
Cadps2
Lancl3
Sbk1
Cpne8
Tnfrsf25
Atg9b
Apold1
